# Nutrient-responsive and DAF-16/FoxO target H1 histone HIL-1 promotes resistance to starvation and bacterial pathogens in *Caenorhabditis elegans*

**DOI:** 10.64898/2026.05.29.728887

**Authors:** Kinsey Fisher, Rojin Chitrakar, L. Ryan Baugh

## Abstract

Insulin/IGF-1 signaling (IIS) mediates metabolic and developmental acclimation to stressful conditions including starvation. The transcription factor DAF-16/FoxO actuates many of the physiological effects of reduced IIS, yet the specific contributions of DAF-16 target genes to stress resistance remain poorly understood. We explore the function of *C. elegans* H1 linker histone HIL-1/H1.0, a DAF-16 target that is upregulated during starvation. The HIL-1 sequence is divergent from the other eight annotated *C. elegans* H1 variants, and the others are not so highly responsive to nutrient availability and DAF-16 activity, suggesting distinct function. Using knock-in reporters, we find that HIL-1 is broadly expressed in nuclei of L1 and dauer larvae during starvation, but that expression is largely undetectable in fed larvae. Disrupting *hil-1* activity through mutation or auxin-inducible degradation led to reduced growth after extended L1 starvation, revealing reduced starvation resistance. RNA-seq of *hil-1* mutants suggested that HIL-1 activates genes involved in the innate immune response, and *hil-1* mutants display compromised survival upon exposure to *Pseudomonas aeruginosa* under reduced IIS. Together these results suggest that DAF-16/FoxO activates transcription of *hil-1* during starvation to promote resistance to starvation and pathogens. We demonstrate conditional regulation of an H1 histone, and we reveal a novel mechanism for how IIS promotes stress resistance by identifying a histone variant that connects nutrient sensing to immunity.

**ARTICLE SUMMARY:** H1 histones promote condensation of chromatin. Like many other organisms, the roundworm *Caenorhabditis elegans* has several H1 histones, but their regulation and function is unclear. We were intrigued by *hil-1/H1.0* because it is dramatically up-regulated during starvation in *daf-16/FoxO*-dependent fashion. Insulin/IGF signaling regulates DAF-16, and DAF-16 is critical to starvation resistance. We used genome editing to create alleles to interrogate *hil-1* expression and function. We show HIL-1 is expressed throughout the animal during starvation, it activates expression of immunity genes, and it promotes resistance to starvation and a bacterial pathogen. This work demonstrates a connection between nutrient sensing and immunity.

## INTRODUCTION

Wild animals must contend with fluctuating, often stressful, environmental conditions, and they must have robust physiological responses to survive. The nematode *Caenorhabditis elegans* consumes ephemeral microbial sources of food in the wild (Schulenburg and Félix 2017). *C. elegans* larvae develop into dauers as an alternative to the third larval stage in response to high population density and limited food (Baugh and Hu 2020), and when larvae hatch in the absence of food they reversibly arrest development in the first larval stage (L1 arrest or L1 diapause) (Baugh 2013). *C. elegans* L1 arrest and recovery is a powerful animal model to study how animals respond to starvation.

The widely conserved insulin/IGF-1 signaling (IIS) pathway coordinates growth, development, and metabolism with nutrient availability. IIS antagonizes FoxO transcription factors, which promote longevity, tumor suppression, DNA repair, and cell cycle arrest in mammals (Carter and Brunet 2007). In humans, hyperactivation of insulin signaling is common in cancer (Belfiore 2007), and dysregulation of insulin signaling is central to diabetes (Boucher et al. 2014). Moreover, IIS plays a critical role in regulation of ageing (Kenyon 2011). The sole known insulin/IGF-1 receptor in *C. elegans,* DAF-2/InsR, activates PI3K/AKT signaling, which antagonizes DAF-16/FoxO to promote stress resistance, developmental arrest, metabolic adaptation, and longevity (Murphy and Hu 2013). *daf-16* promotes developmental arrest and survival in starved L1 larvae (Baugh and Sternberg 2006; Muñoz and Riddle 2003), contributing to the starvation response by regulating thousands of genes involved in metabolism, homeostasis, and immunity (Fisher et al. 2026; Hibshman et al. 2017; Kaplan et al. 2015). Despite IIS being so well studied, the function of most DAF-16 targets in mediating the starvation response remains unclear.

H1 histones bind linker DNA and pull nucleosomes together to modulate chromatin compaction and gene expression (Di Liegro et al. 2018). They are highly conserved and have a tripartite structure consisting of two unstructured and variable domains, a short N-terminal domain (NTD) and a long lysine-rich C-terminal domain (CTD), as well as a central, evolutionarily conserved globular domain (GD) with a winged-helix structure (Hartman et al. 1977). The canonical function of the GD is to bind to the nucleosome, while the CTD binds to linker DNA through electrostatic interactions. The role of the NTD is not as well defined, but it is thought to contribute to nucleosome binding (Sridhar et al. 2020) and protein-protein interactions (Kalashnikova et al. 2013).

H1 variants have different expression patterns, with some being found exclusively in certain cell types such as testes or terminally differentiated cells, or at specific phases of the cell cycle (Harshman et al. 2013). They also have different binding affinities for chromatin (Catez et al. 2006; Orrego et al. 2007). Chromatin binding affinity of H1 variants is primarily driven by the CTD (Hendzel et al. 2004), and post-translational modifications of the CTD and NTD direct H1 interactions with DNA and other proteins (Kalashnikova et al. 2016). Nutrient-dependent regulation of H1 histone expression has not been reported.

The *C. elegans* genome encodes nine annotated H1 variants. This gene family was identified in the 1990s (Sanicola et al. 1990), but understanding of the physiological function of *C. elegans* H1 variants is limited, and the extent to which the H1 genes have overlapping vs. specific function is unclear. The H1 variant-encoding gene *hil-1* is a direct target of DAF-16/FoxO (Schuster et al. 2010; Tepper et al. 2013), and it is dramatically up-regulated during L1 arrest in *daf-16*-dependent fashion (Baugh et al. 2009; Fisher et al. 2026). In addition, HIL-1 (previously known as H1.X) has a divergent protein sequence compared to other *C. elegans* H1 histones, with hydrophobic residues on its terminal domains (Jedrusik and Schulze 2001). Surprisingly, HIL-1 is reported to function in the cytoplasm to control muscle growth and function (Jedrusik et al. 2002). However, *hil-1* has very low expression during development compared to starvation (Baugh et al. 2009; Fisher et al. 2026), and its role in starvation has yet to be investigated.

We characterized the role *hil-1* plays in L1 arrest and recovery with phenotypic, reporter gene, and transcriptome analyses. We show that nutrient availability and IIS govern *hil-1* expression, distinguishing it from other H1 histones. We report a novel function for *hil-1*, linking nutrient availability to immunity, and we identify a bacterial pathogen-sensitive phenotype for *hil-1* mutants under reduced IIS. This work uncovers a novel mechanism for how DAF-16/FoxO promotes stress resistance through its target genes, and it provides an example of an H1 variant that has evolved conditional regulation and distinct function.

## MATERIALS AND METHODS

### Bacterial strains and preparation

#### Preparation of OP50 for solid media

One colony of *E. coli* strain OP50 was added to 100 mL of LB and grown overnight at room temperature then stored at 4°C. Four drops (∼400 μL) or one drop (∼100 μL) of OP50 were added to 10 cm and 6 cm plates of nematode growth medium (NGM), respectively, and plates were incubated at room temperature for two days before using them for culturing worms and recovering them from starvation.

#### Preparation of HB101 for liquid culture

One colony of *E. coli* strain HB101 was added to a 5 mL starter culture of LB with 50 μg/mL streptomycin and grown overnight at 37°C in a shaking incubator. The starter cultures were then added to a 1 L culture of TB with streptomycin and grown for 24 hours at 37°C. The bacteria were centrifuged for 10 min at 4,000 rpm to form a pellet. After being weighed, the bacterial pellet was resuspended in S-complete to create a 10x (250 mg/mL) stock that was stored at 4°C. Further dilutions with S-complete were used to create the dilutions for each condition in this experiment. All “fed” cultures, except for those in the dilution series (Figure 2f), included 25 mg/mL HB101 (Hibshman et al. 2021).

#### Preparation of *P. aeruginosa* PA14 fast-killing plates

One colony of *P. aeruginosa* strain PA14 was picked into 3 mL of LB in a 15 mL conical tube and grown at 37°C in a shaking incubator for 12-16 hr. 15 μL of culture were spread onto 6 cm Peptone-Glucose-Sorbitol (PGS) agar plates to cover the entire plate and grown overnight at 37°C and then an additional 24 hr at room temperature (Kirienko et al. 2014).

#### Preparation of RNAi plates

Frozen glycerol stocks were streaked on to LB plates with carbenicillin (100 µg/mL) and tetracycline (12.5 µg/mL) and grown overnight at 37°C. A single colony was picked into 1 mL LB with 50 µg/mL carbenicillin and 12.5 µg/mL tetracycline and grown overnight at 37°C in a shaking incubator. 100 µL of this culture was used to inoculate 5 mL of TB with 50 mg/mL carbenicillin and grown overnight at 37°C in a shaking incubator. Cultures were then centrifuged at 4000 rpm for 10 min to form a pellet. After being weighed, the bacteria were then resuspended in S-complete with 20% glycerol at a 4:1 (S-complete:bacteria) ratio by mass. 100 µL aliquots were frozen and stored at - 80°C. 40 µL from one of these aliquots was spread onto 6 cm NGM plates with 50 mg/mL carbenicillin and 1 mM IPTG, and plates were cultured overnight at room temperature before plating worms for RNAi treatment.

#### *C. elegans* maintenance and strains

Strains used in this study are described in Table 1. *C. elegans* strains were maintained at 20°C on NGM plates seeded with *E. coli* OP50.

**Table 1.** Worm strains used in this study. Each worm strain used in this study, with details for full genotype, function, and origin.

| Strain | Genotype | Description | Source |
| --- | --- | --- | --- |
| N2 | Wild type | Sternberg N2 | Paul Sternberg – California Institute of Technology |
| CB1370 | *daf-2(e1370)* | *daf-2* loss-of-function allele | CGC |
| LRB437 | *hil-1(gk229)* | Deletion (236 bp) | CGC<br>Backcrossed 4x to N2 |
| LRB438 | *hil-1(tm1442)* | Deletion (960 bp) | Shohei Mitani - Tokyo Women's Medical University School of Medicine<br>Backcrossed 4x to N2 |
| LRB439 | *daf-2(e1370); hil-1(gk229)* | *hil-1(gk229)* crossed to *daf-2(e1370)* | Generated in this work |
| LRB440 | *daf-2(e1370); hil-1(tm1442)* | *hil-1(tm1442)* crossed to *daf-2(e1370)* | Generated in this work |
| LRB678 | *hil-1(syb9165)* | Full deletion of *hil-1* using CRISPR | Created by SUNY Biotech and gifted by Sevinc Ercan's lab<br>Backcrossed 4x to N2 |
| LRB696 | *daf-2(e1370); hil-1(syb9165)* | *hil-1(syb9165)* crossed to *daf-2(e1370)* | Generated in this work |
| PHX5748 | *hil-1(syb5748 [GFP::linker::AID::linker::3xFLAG::linker::hil-1])* | GFP, AID and 3xFLAG tagged to endogenous *hil-1* N-terminus with CRISPR | Generated in this work by SunyBiotech |
| PHX5764 | *hil-1(syb5764 [hil-1::linker::GFP::linker::AID::linker::3xFLAG])* | GFP, AID and 3xFLAG tagged to endogenous *hil-1* C-terminus with CRISPR | Generated in this work by SunyBiotech |
| LRB556 | *daf-2(e1370); hil-1(syb5748)* | *hil-1(syb5748)*<br>crossed to *daf-2(e1370)* | Generated in this work |
| LRB546 | *daf-2(e1370); hil-1(syb5764)* | *hil-1(syb5764)*<br>crossed to *daf-2(e1370)* | Generated in this work |
| LRB545 | *daf-16(mu86); hil-1(syb5748)* | *hil-1(syb5748)*<br>crossed to *daf-16(mu86)* | Generated in this work |
| LRB547 | *daf-16(mu86); hil-1(syb5764)* | *hil-1(syb5764)*<br>crossed to *daf-16(mu86)* | Generated in this work |
| LRB543 | *KeaSi10[rpl-28p::TIR1::mRuby::unc-54 3' UTR+(br-unc-119(H))]; hil-1(syb5748)* | *hil-1(syb5748)*<br>with ubiquitous TIR1 expression | Generated in this work |
| LRB544 | *KeaSi10[rpl-28p::TIR1::mRuby::unc-54 3' UTR+(br-unc-119(H))]; hil-1(syb5764)* | *hil-1(syb5764)*<br>with ubiquitous TIR1 expression | Generated in this work |

### Preparation of strains before bleach

Seven L4 larvae were picked onto 10 cm plates seeded with OP50 and cultured at 20°C for 96 hr (or 120 hr for strains with *daf-2(e1370)* to account for growth delay). Plates were washed with S-basal and treated with hypochlorite solution to obtain sterile, developmentally synchronized embryos (Hibshman et al. 2021).

### Phylogenetic analysis

Full-length protein sequences were obtained from UniProt (Consortium 2024b) and downloaded in FASTA format. Multiple-sequence alignments were done for all histone sequences across the five model organisms (Figure 1c) using EMBL-EBI’s ClustalOmega (Sievers et al. 2011), and alignments were output in the “NEXUS” format. The unrooted, distance-based phylogenetic tree was generated by ClustalOmega’s default methods from the multiple sequence alignment.

**Figure 1.**
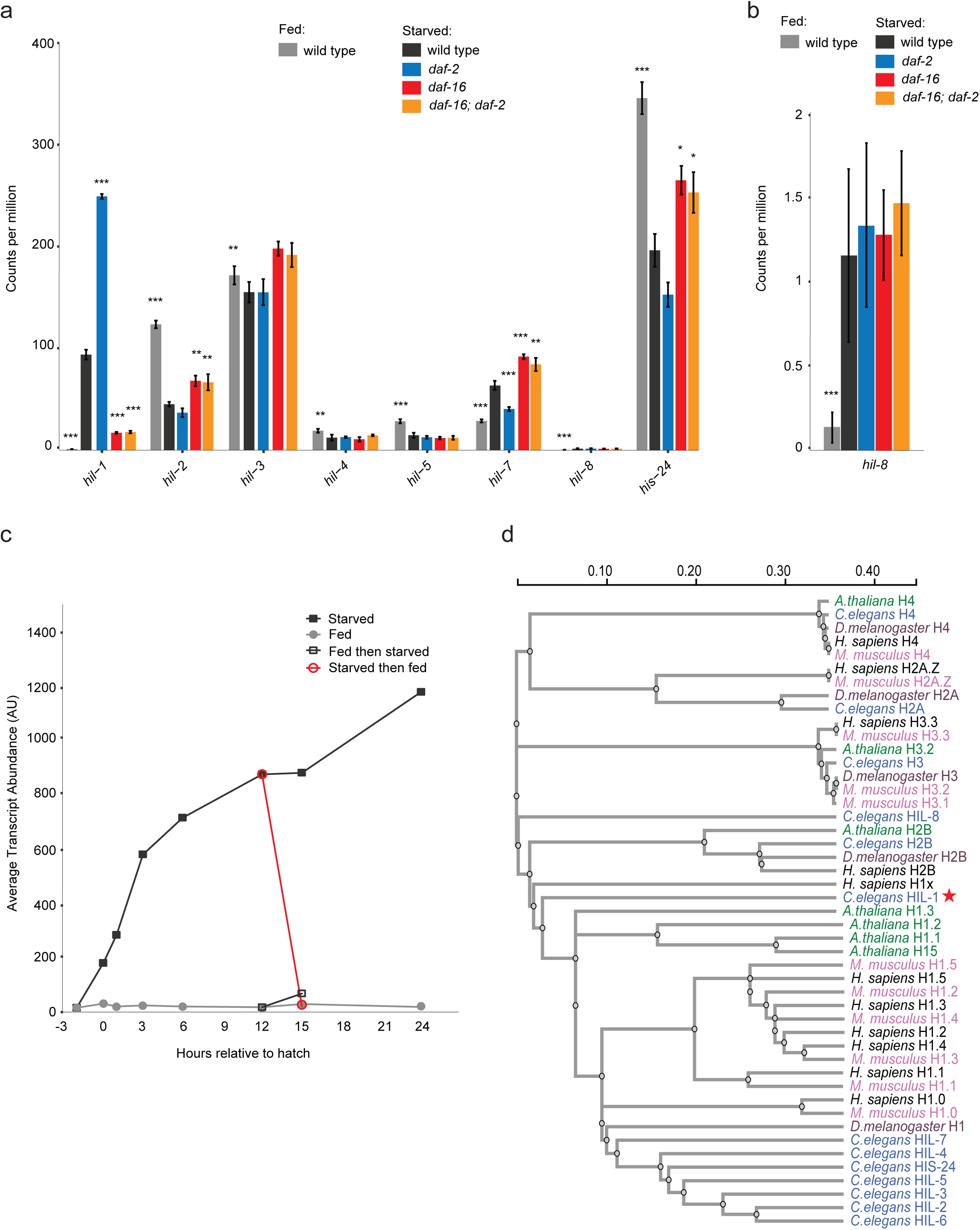
HIL-1 regulation and sequence are distinct from other *C. elegans* H1 histones. a, b) Transcript abundance (counts per million) from RNA-seq ∼12 hours after hatching for eight of nine detected H1 variants in fed and starved wild-type worms as well as starved *daf-16(mgDf47)*, *daf-2(e1370)*, and *daf-16(mgDf47); daf-2(e1370)* mutants (Fisher et al. 2026). *P < 0.05, **P < 0.01, ***P < 0.001; 3-4 biological replicates; error bars reflect standard deviation. (FDR from exact test against wild type starved). b) *hil-8* transcript abundance with a smaller scale to visualize differences across conditions. c) Wild-type *hil-1* expression in microarray timeseries of fed and starved L1 larvae (Baugh et al. 2009). Fed and starved larvae were hatched with or without food and sampled over time; fed then starved larvae were hatched with food, were starved 12 hr later, and were sampled 3 hr after that (red open circles); and starved then fed larvae were hatched without food, were fed 12 hr later, and were sampled 3 hr after that (black open squares). Interaction P-value from a two-way ANOVA < 10^-16^; three biological replicates. See Fig. S1 for additional recovery data. d) Phylogenetic tree of histone protein sequences across five popular multicellular organisms. *C. elegans* HIL-1 is indicated with a red star.

### *C. elegans* liquid cultures

After hypochlorite treatment (see above), 5,000 embryos were placed into 5 ml of S-basal (starved) or S-complete with HB101 (fed) in 16 mm glass test tubes at 20°C in the dark on a tissue culture roller drum at ∼30 rpm. Embryos hatched and arrested as L1 larvae in starved cultures. Aliquots from these cultures were used for starvation resistance assays and reporter imaging.

### High-magnification imaging of GFP in L1 larvae

To obtain 1000x images of the *hil-1* GFP reporters, 1,000 L1 larvae were pulled from the liquid culture and rinsed 3x with S-basal (starved) or S-complete (fed) and resuspended in 10 mM levamisole in a 1.5 mL Eppendorf tube. Worms were spun at 3,000 rpm for 1 min to pellet and transferred to 4% noble agar pads on a microscope slide. Worms were imaged at 1000X total magnification on a Zeiss AxioImager compound microscope using Nomarski microscopy or a GFP filter. Exposure times were equivalent between fed and starved samples within a strain. GFP and Nomarski images were merged using Fiji (Schindelin et al. 2012).

### High-magnification imaging of GFP in dauer larvae

To obtain 1000x images of the *hil-1* GFP reporters in dauer and L3 larvae, 20 L3 or dauer larvae were picked from a fed or recently starved plate, respectively, and placed onto a 4% noble agar pad with 3 µL of 10 mM levamisole. Imaging was performed the same as above, with the exception that multiple images were taken in each channel to capture the entire body of the worm. Images were stitched together using automated stitching in Fiji (Preibisch et al. 2009), and black edges from stitching were removed using content-aware filling on the background in Photoshop version 27.2.0.

### High-magnification imaging of GFP after 24 and 48 hours of feeding

After 24 and 48 hr of continuous feeding from hatching or from recovery of larvae that had been starved 12 hr (Fig. 2e, Fig. S3), 500 µL aliquots were pulled from the cultures and rinsed, paralyzed, and plated onto 4% noble agar pads as described above. High exposure times were used to capture relatively low GFP expression, with consistent exposure times across all conditions and timepoints within each strain. Images were stitched and prepared for display as described above.

**Figure 2.**
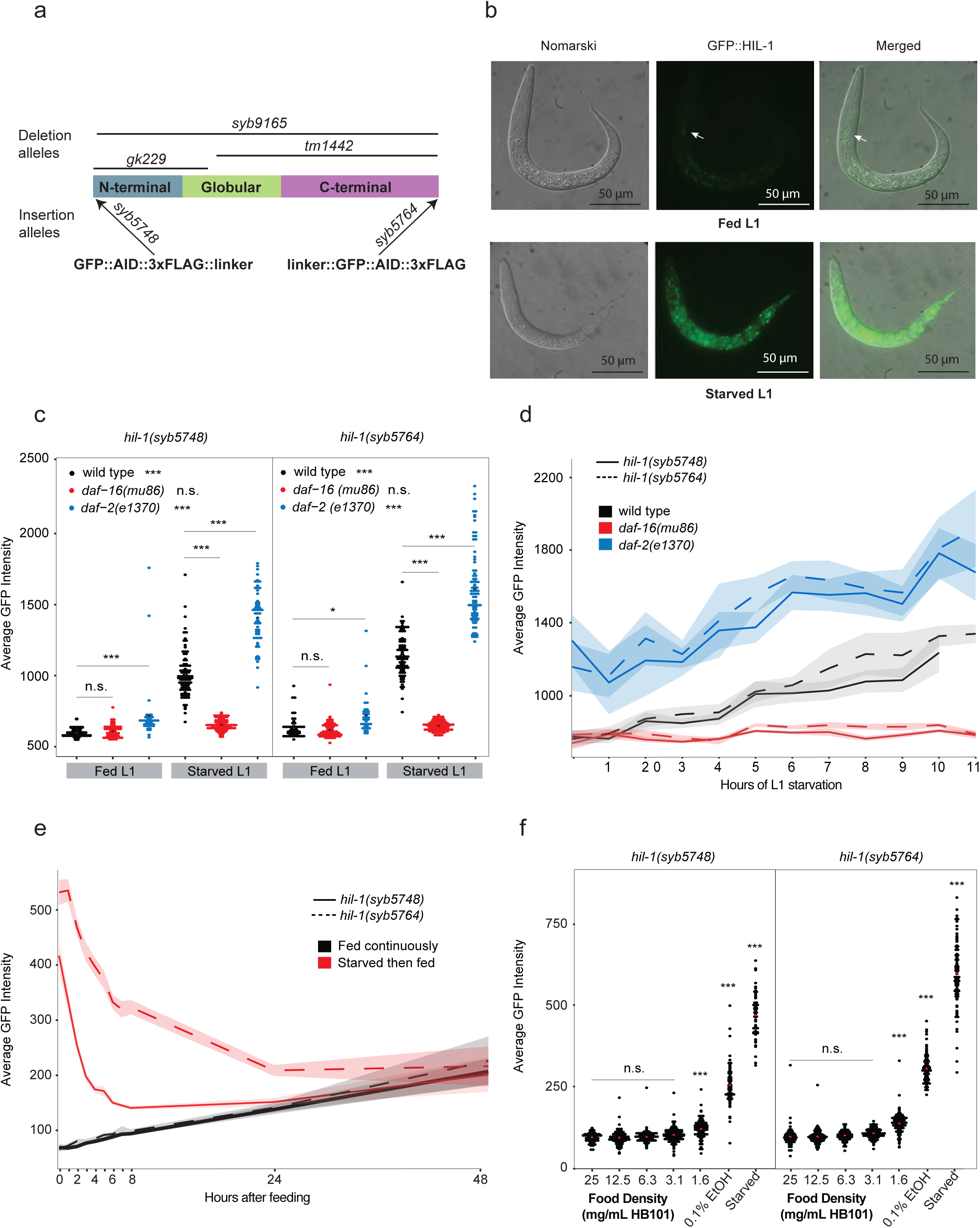
HIL-1 is expressed broadly in the nuclei of starved larvae under the control of DAF-16/FoxO. a) Diagram of HIL-1 protein structure and five different *hil-1* alleles used in this study. b) Nomarski and GFP images of *hil-1(syb5748)* in fed (25 mg/ml HB101) and starved L1 larvae ∼12 hr after hatching at 1000x. Identical exposure times were used for fed and starved larvae. Arrows point to relatively faint pharyngeal GFP expression in fed larvae. See Fig. S2a for *hil-1(syb5764)*. See Fig. S3 for longer exposure times of fed larvae. c) Average background-corrected GFP intensity of *hil-1* reporters in fed and starved larvae ∼12 hr after hatching in wild type, *daf-16(mu86)* and *daf-2(e1370)*. * P <0.05, *** P < 0.001, n.s. not significant (one-way ANOVA and *post-hoc* Tukey test; legend asterisks indicate fed vs. starved within genotype, and asterisks over horizontal lines indicate mutant vs. wild type within a condition). d) Average background-corrected GFP intensity of *hil-1* reporters over time (every hour) after hatching without food (starvation) in wild type, *daf-16(mu86)* and *daf-2(e1370).* e) Average background-corrected GFP intensity of *hil-1* reporters over time (time points indicated) during recovery from ∼12 hr L1 starvation (starved then fed) and over time after hatching with food (continuously fed). Increasing signal over time in continuously fed larvae is due to increasing autofluorescence and not GFP expression. See also Fig. S3. d-e) Shading reflects the 95% confidence interval; 30-80 animals per data point. Tick marks on the x-axis indicate time points where data was collected. f) Average background-corrected GFP intensity of *hil-1* reporters ∼12 hours after hatching without food, without food but supplemented with 0.1% ethanol (EtOH), or in variable densities of food (HB101 densities indicated). *** < 0.001 compared to all other conditions; n.s. not significant from others in group indicated by line (one-way ANOVA and post-hoc Tukey test); two biological replicates. Refer to Fig. S2.

### Quantification of GFP using automated imaging

Worms were collected from liquid cultures, pelleted, and rinsed as described above, then transferred to a 96-well plate with 50 μM sodium azide. Plates were imaged using an ImageXpress® Nano automated imager at 100X total magnification under transmitted light and GFP channels using the same exposure time for all samples within an experiment. Objects were filtered by size (5 µm - 40 µm width and 10000 - 75000 pixels), and background intensity was subtracted. Detected objects were manually screened to remove any overlapping, segmented or unfocused worms and debris. Statistical differences between treatment groups were determined using one-way ANOVA and post-hoc Tukey tests. Plots were generated using ggplot2 in R.

### Timeseries imaging

For the starvation timeseries (Fig. 2d), 380 µL (∼380 worms) was pulled from each starvation culture and added directly to a 96-well plate with 50 μM sodium azide for imaging each hour after hatching (∼12 hours after hypochlorite treatment equals 0 hr starvation). For the starved then fed timeseries (Fig. 2e), starvation cultures were established as previously described. 24 hr after hypochlorite treatment (∼12 hr post-hatch), images were acquired to define time 0 (before feeding starved cultures), after which starved cultures were fed HB101 at a final density of 25 mg/mL. For the continuously fed timeseries (Fig. 2e), cultures were set up by hypochlorite treatment at a 12 hr offset from starved cultures, such that larvae hatched directly into 25 mg/mL HB101 and were developmentally synchronized with the starved then fed cultures. Worms were imaged every hour for the first 0–8 hr of feeding, and again at 24 and 48 hr. High-magnification images were acquired at 12, 24, and 48 hr post-feeding as described above to visualize relatively low GFP expression in fed larvae. The same culture was used for the full timeseries for each genotype and condition. Automated images using Molecular Devices ImageXpress® Nano was used for imaging and analysis as described above for all three timeseries.

### Starvation survival

For each day of L1 arrest, ∼100 worms (100-150 μL) were plated off the the lawn of a 6 cm NGM plate seeded with a small drop (∼100 μL) of *E. coli* OP50 in the center and placed at 20°C, and the total number of worms plated (dead or alive) was counted (number plated). Two days after plating, the number of worms that were crawling on the lawn were counted (survivors). Survival was determined by dividing the number of survivors by the number plated. 3-4 biological replicates were used for each genotype. A quasi-binomial generalized linear model was used to fit the proportion of survival to the duration of L1 arrest for each replicate. Median survival was determined by the model for each replicate. Medians of each group of replicates per genotype were tested for homogeneity of variance using Bartlett’s test. Two-tailed, unpaired t-tests were performed on the medians for each genotype with variance pooled across genotypes (assuming homogeneity). Plots were generated using ggplot2 in R.

### Auxin-inducible degradation (AID)

A 400 mM master stock of synthetic, water-soluble auxin analog (1-naphthaleneacetic acid, potassium salt [K-NAA] (Martinez et al. 2020)) was prepared in water and stored in the dark at −20°C. Working stocks of 40 mM auxin were prepared by diluting auxin in water and storing in the dark at −20°C. To degrade HIL-1 during starvation, 100 μM auxin was added to the starvation culture from the working stock after adding embryos from the hypochlorite treatment. Degradation of HIL-1 was confirmed by dramatic loss of GFP signal when observed at 1000x on a Zeiss AxioImager compound microscope (Fig. S4c, d). Relatively weak GFP signal remained after addition of auxin, indicating strong but incomplete knock-down of HIL-1.

### Length measurement following recovery from L1 starvation

At day 1 and day 8 of starvation, ∼1000 worms (1 mL and 1.5 mL aliquots of the starvation culture for day 1 and day 8, respectively) were placed into 15 mL conical tubes and pelleted at 3,000 rpm for 1 min. L1 worm pellets were plated onto 10 cm NGM plates seeded with a lawn of *E. coli* OP50 and placed at 20°C. After 48 hours, worms were rinsed off the plates using S-basal and placed into 15 mL conical tubes and pelleted at 3,000 rpm for 1 min. Worm pellets were then placed onto an unseeded 10 cm plate and allowed to dry for ∼10 min. Images were taken on a ZeissDiscovery V20 stereomicroscope at 20x (day 1) or 40x (day 8) magnification. Worm length was determined using the WormSizer FIJI plugin as previously described (Moore et al. 2013). 3-4 biological replicates were imaged and analyzed for each experiment. Using the “nlme” package in R, a two-factor model was used to assess genotype- or condition-dependent effects of starvation on length, with interaction P-values determined by fitting a linear mixed-effect model with fixed effects for genotype or condition (addition of auxin) and duration of starvation with a random effect of replicate. Pairwise comparisons between strains at day 1 or day 8 were calculated using “emmeans” on the linear model in R. Plots were generated using ggplot2 in R.

### Length measurement without starvation

50 young gravid adults were picked onto a 10 cm NGM plate seeded with a lawn of *E. coli* OP50 for each strain and left on the plate for 1 hour at 20°C. Adults were then removed by picking and the cultures were left for 60 hr at 20°C. After 60 hr, the worms were washed, replated on unseeded 10 cm NGM plates, imaged, and analyzed as described above.

### Sample preparation and collection for bulk RNA-seq

Worms were prepared and treated with hypochlorite solution as described above. 20,000 embryos were placed in individual cultures of 20 mL S-basal in 50 mL flasks. All flasks were incubated at 20°C and 180 rpm for 24 hours. With these conditions, it takes approximately 12 hours for embryos to hatch following hypochlorite treatment. Samples were then washed 3x with S-basal before being pelleted and snap frozen with liquid nitrogen. Samples were stored at -80°C until RNA isolation and library preparation. Four biological replicates were collected for each genotype.

### RNA isolation and library preparation

RNA was isolated from samples with TRIzol Reagent (Invitrogen#15596026) per manufacturer’s instructions, except 100 uL of acid-washed sand (Sigma-Aldrich #27439) was added to each sample at the beginning of the extraction protocol to aid with homogenization. RNA was eluted in nuclease-free water and stored at -80°C until further use. Libraries were prepared for sequencing using the NEBNext Ultra II RNA Library Prep Kit for Illumina (New England Biolabs # E7775) starting with 1 ug of total RNA per sample as input and seven cycles of PCR. Individually barcoded libraries were pooled and sequenced on a single lane of the NovaSeq X Plus 10B flowcell to obtain 50 bp paired-end reads.

### Read mapping and differential expression analysis

Read mapping was performed as described in Fisher et al. 2026. In short, Bowtie was used to map reads to version WS273 of the genome (Langmead et al. 2009). HTSeq was used to count mapped reads per gene model (Putri et al. 2022). Count data was restricted to include only protein-coding genes. An ExactTest in EdgeR was used for differential expression analysis (Chen et al. 2025) on genes with counts per million (CPM) > 1 in at least four libraries across all samples. Differential expression analysis plot was generated using “plotSmear” function in R.

### Cluster analysis of *hil-1* targets on chromosomes

Detected genes from RNA-seq were mapped to *C. elegans* genomic coordinates using biomaRt (Durinck et al. 2009) and GenomicRanges (Lawrence et al. 2013). Spatial clustering of target genes (significantly up or down by *hil-1* in RNA-seq) was assessed using regioneR permutation testing (Gel et al. 2016) (1,000 permutations), in which random gene sets of equal size were sampled from the background (all detected genes in RNA-seq) and evaluated using mean nearest-neighbor distance method.

### Gene Ontology-term Analysis

Gene Ontology (GO) enrichment analysis for “biological process” (Ashburner et al. 2000; Consortium et al. 2023; Thomas et al. 2022) was preformed separately for genes activated and repressed by *hil-1* (FDR < 0.1) using GOrilla (Eden et al. 2009). Background list was all protein-coding genes detected in the experiment (13,068 genes). Visualization was provided by GOrilla (Eden et al. 2009). “Driver terms” in gene ontology are those that summarize many related, significantly enriched terms. gProfiler (Kolberg et al. 2023) was used to determine the driver terms for genes activated by *hil-1*, which resulted in only one driver term, “defense response to other organism”. This term was used for identification of *hil-1* targets to use in the candidate screen.

### Estimation of transcription factor activity using CelEsT

We used CelEsT v1.1 (Perez 2024) to infer changes in transcription factor activity based on bulk RNA-seq results comparing *hil-1* deletion mutants and wild type. We used replicate-level count data as input, and CelEsT performed differential expression analysis with DESeq2 (Love et al. 2014) and used those results to assess changes in transcription factor activity based on its gene regulatory network model. The output table containing Z-scores and P-values for each transcription factor was downloaded (Table 2; Sup. File 1) and ggplot2 was used to visualize the results.

**Table 2.**
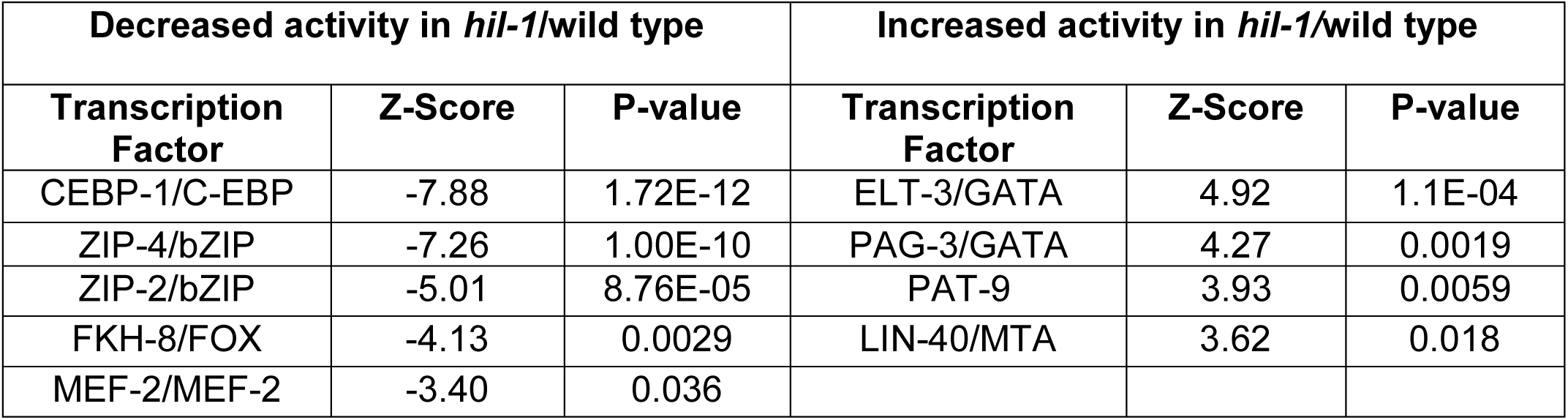
*hil-1*-dependent changes in transcription factor activity inferred with CelEsT. CelEsT P-values were adjusted for multiple testing (Benjamini-Hochberg), and all transcription factors with significantly altered activity are included here.

### Comparisons of differentially expressed genes to other datasets

We compared our *hil-1-*regulated genes to four different published datasets: 1) Genes that were up- or down-regulated on *P.aeruginosa* PA14 (Padj < 0.05; 890 and 803 genes, respectively) (Fletcher et al. 2019) 2) CEBP-1 ChIP-seq targets (212 genes) (Kim et al. 2016), 3) genes that are up- or down-regulated in *nipi-3(fr4)* compared to wild type on PA14 (FDR < 0.05 and Log_2_ fold-change > 1 or < -1; 282 and 70 genes, respectively) (McEwan et al. 2016), and genes that are differentially expressed on PA14 in a *pmk-1*-dependent fashion (Fletcher et al. 2019). Geometrically accurate Venn diagrams were created using the “eulerr” package in R and statistical analyses of overlaps between datasets were conducted using hypergeometric tests with corrections for multiple testing using the Bonferroni method.

### *C. elegans* survival on *P. aeruginosa* PA14 fast-killing plates

Embryos were synchronized using hypochlorite treatment and plated directly onto OP50 or RNAi plates (for the candidate screen). After 48 hours (wild-type background) or 56 hours (*daf-2(e1370)* background), ∼50-100 L4 worms were transferred from OP50 or RNAi plates onto fast-killing PA14 plates. For the long-lived *daf-2(e1370)* mutants, surviving worms were transferred to a fresh fast-killing plate every other day until all worms were dead. Survival was scored by movement after being gently prodded with a worm pick. Dead worms were removed from the plate and all worms that escaped the lawn by crawling up the side of the plate were censored. Survival was scored every 2 hours for the first 6 hours, then every 24 hours until no living worms remained. Four biological replicates were collected for each strain or condition in each experiment. Statistical analysis between strains or RNAi treatments was preformed using a log-rank test in Oasis 2 (Han et al. 2016). Survival step-plots were generated using ggplot2 in R.

### C. elegans lifespan on E. coli OP50

30 L4 worms were picked onto 6 cm plates seeded with one drop (∼100 uL) of *E. coli* OP50 for each strain. Each day, the number of dead worms was scored and surviving worms were transferred to a fresh OP50 plate every other day until all worms were dead. Survival was scored by movement after being gently prodded with a worm pick. Dead worms were removed from the plate and all worms that escaped the lawn by crawling up the side of the plate were censored. Three biological replicates were collected for each strain or condition. Statistical analysis between strains was preformed using a log-rank test in Oasis 2. Survival step-plots were generated using ggplot2 in R.

### Determination of genes used in candidate screen

The three criteria used to identify candidate genes is described in Results. Candidates for which RNAi bacteria was not available were not tested. These restrictions (plus addition of *nipi-3*; see Results) resulted in 12 candidate genes for the screen, in addition to empty vector and *hil-1* RNAi controls.

## RESULTS

### *hil-1* transcriptional regulation and protein sequence are unique among *C. elegans* H1 histones

We previously used bulk RNA-seq to characterize mRNA expression in fed and starved wild-type L1 larvae as well as a *daf-16/FoxO* null mutant (*mgDf47*) (Lee et al. 2001), a class II loss-of-function *daf-2/InsR* mutant (*e1370*) (Gems et al. 1998), and a *daf-16(mgDf47); daf-2(e1370)* double mutant in starved L1 larvae (Fisher et al. 2026). We interrogated expression of the *C. elegans* H1 histone gene family with this dataset (Fig. 1a, b). All H1 histone genes were differentially expressed in response to nutrient availability (fed vs. starved) in wild-type animals. *hil-2, hil-3, hil-4, hil-5,* and *his-24* were expressed at significantly higher levels in fed larvae, and *hil-1, hil-7*, and *hil-8* were expressed at higher levels in starved larvae (Fig. 1a, b). *hil-1* stands out among H1 histones with its large dynamic range of expression: It is barely detected in fed larvae (counts per million = ∼1), and it is upregulated ∼85-fold during starvation. Results from another study suggest that *hil-1* upregulation continues for at least 24 hr into L1 starvation while it was not detected throughout L1 and L2 development in fed larvae (Fig. 1c)(Baugh et al. 2009). These results show that *hil-1* is unique among *C. elegans* H1 histones in its transcriptional responsiveness to nutrient availability.

Four of the annotated H1 genes, *hil-1, hil-2, hil-7* and *his-24,* were differentially expressed in *daf-16* and *daf-16; daf-2* mutants compared to wild type during starvation (Fig. 1a), suggesting DAF-16 regulates their transcription downstream of IIS. Notably, *hil-1* was the only variant downregulated in the *daf-16* mutants. *hil-1* is the only H1 variant that is directly bound by DAF-16, and DAF-16 is believed to function as an activator of its direct targets (Schuster et al. 2010; Tepper et al. 2013). These observations suggest that DAF-16 directly activates transcription of *hil-1* during starvation or other conditions with low IIS. This putative regulatory mechanism further distinguishes *hil-1* from the other *C. elegans* H1 genes.

The HIL-1 sequence diverges from other H1 histones, including those of *C. elegans*, especially in the NTD and CTD (Jedrusik and Schulze 2001). We performed phylogenetic analysis of histone protein sequences across five diverse, multicelluar organisms (Fig. 1d). HIL-1, -2, -3, -4, -5, and -6 as well as HIS-24 are each described as orthologs of human H1.0 on the Alliance of Genome Resources (Consortium 2024a). H1 histones from all five eukaryotes clustered together, but HIL-1 is clearly distinct from the other *C. elegans* H1 histones as well as those from human, mouse, *Drosophila*, and *Arabidopsis*. These results suggest an expansion of the H1.0 common ancestor into a family of paralogs during the evolution of *C. elegans*, with HIL-1 diverging from the others. Notably, although HIL-8 is classified as an “H1-like” protein on the Alliance of Genome Resources (Consortium 2024a), it is more similar to H2B than H1 histones. Together with its unique transcriptional regulation, divergence of the HIL-1 protein sequence from other H1 histones suggests it may have distinct function.

### Widespread nuclear expression of HIL-1 in starved larvae under the control of daf-16/FoxO

We used CRISPR-Cas9 to make endogenous GFP reporter genes for HIL-1. We generated two alleles, one with a knock-in on the N-terminus (*syb5748)* and one on the C-terminus (*syb5764*). In addition to GFP, the knock-ins include an auxin-inducible degron (AID), 3x FLAG tag, and flexible linkers (Fig. 2a). When fed for ∼12 hr after hatching, GFP was invisible except for faint signal in the isthmus of the pharynx (Fig. 2b, S2a, S3a, b). In stark contrast, robust expression was evident throughout the animal in worms that had been starved for ∼12 hr after hatching (Fig. 2b, S2a), though we cannot definitively conclude that it is expressed in every cell. HIL-1 was also expressed in several major tissues in dauer larvae (Fig. S2b, c). Notably, expression was largely nuclear, consistent with canonical function as an H1 histone.

We quantified whole-worm GFP expression in fed and starved larvae ∼12 hr after hatching in wild-type, a *daf-16* null mutant, and *daf-2(e1370)*. HIL-1 expression was significantly upregulated during starvation in wild-type worms for both reporter alleles (Fig. 2c). Up-regulation in response to starvation was increased by mutation of *daf-2/InsR*, and disruption of *daf-2* caused relatively modest upregulation in fed larvae. Loss of *daf-16/FoxO* abrogated upregulation in response to starvation, and *daf-16* was required for upregulation by *daf-2(e1370)* in fed and starved larvae (Fig. 2c). Timeseries analysis revealed similar patterns of regulation: HIL-1 expression was immediately upregulated upon hatching in the absence of food and continued increasing for at least 11 hours, upregulation was amplified by *daf-2(e1370)*, and there was no upregulation with loss of *daf-16* (Fig. 2d). HIL-1 reporter gene analysis using two different knock-in alleles corroborates transcriptome analysis (Fig. 1a, c), demonstrating dramatic nutrient-dependent regulation under the control of IIS and DAF-16/FoxO.

*hil-1* expression is downregulated as dramatically by feeding as it is upregulated by starvation. Published transcriptome analysis suggests that *hil-1* transcript abundance is rapidly decreased in response to feeding L1 larvae after being starved for ∼12 hr (Fig. 1c) (Baugh et al. 2009), with a return to baseline levels within just 1 hr of feeding (Fig. S1) (Maxwell et al. 2012). Our HIL-1 reporters were also downregulated upon feeding starved larvae (Fig. 2e), though not as quickly as seen for *hil-1* mRNA (Fig. 1c, S1). *hil-1(syb5764)* is brighter than *syb5748* (Fig. 2c, d, f), and *syb5764* GFP signal decays slower during recovery from starvation than *syb5748* (Fig. 2e, S3). Residual GFP expression during recovery was most prominent in the isthmus of the pharynx, as seen in continuously fed larvae (Fig. S3a, b). Whole-animal, average expression levels converge with those of continuously fed larvae (*i.e.*, largely autofluorescence) within 48 hours for both alleles (Fig. 2e), but a small fraction of *hil-1(syb5764)* larvae have substantial residual GFP 48 hours after feeding in the pharynx and elsewhere (Fig. S3d). These results suggest that *hil-1* transcription, transcript stability, and likely protein stability are under strong negative regulation in fed larvae.

We mostly contrast starvation with replete conditions, but IIS activity presumably varies continuously, and growth, reproduction, and longevity are tuned to variation in nutrient availability. We subjected our HIL-1 reporter strains to a two-fold dilution series of bacterial food for ∼12 hr after hatching and quantified GFP intensity. We included replete (25 mg/mL *E. coli* HB101), dietary-restriction (3.1 mg/ml), and dauer-forming (1.6 mg/mL HB101) conditions (Hibshman et al. 2021). We also included starvation and starvation supplemented with 0.1% ethanol. Ethanol serves as a carbon source supporting intermediary metabolism and extending starvation survival without promoting postembryonic development (Castro et al. 2012). There were no significant changes in HIL-1 expression across the higher food densities (25 mg/mL to 3.1 mg/mL), but both reporters showed a modest increase in expression with 1.6 mg/mL compared to replete conditions (Fig. 2f). Ethanol alone further increased HIL-1 expression, but not to the same extent as complete starvation. These results reveal dynamic regulation of HIL-1 expression in response to nutrient availability, suggesting HIL-1 could tune physiology to different levels of low nutrient availability and that HIL-1 expression could be used as a quantitative indicator of nutritional status.

### *hil-1* promotes L1 starvation resistance

Given striking upregulation of HIL-1 throughout the animal during starvation, we wanted to know whether it contributes to starvation resistance. We assessed starvation resistance using two partial deletion alleles of *hil-1*, one in-frame deletion that removes the NTD and part of the GD (*gk229*) and another in-frame deletion that removes part of the GD and the entire CTD (*tm1442*), as well as a full deletion (*syb9165*), which is a null allele (Fig. 2a). We initially analyzed L1 starvation survival, and although all three alleles died modestly faster than wild type, none of these effects were significant (Fig. S4a, b). However, recovery from starvation is as important to fitness as survival, and starvation in a variety of conditions and genotypes can affect recovery with no apparent effect on survival (Baugh and Hu 2020). We counted total brood size to analyze reproduction and used image analysis to analyze worm length following 48 hr recovery from 1 day (control condition for synchronization) or 8 days of L1 starvation (extended starvation). We used a two-factor model to quantify the starvation-dependent effects on reproduction and growth, with the interaction P-value reflecting significance of differences in the reaction norm between 1 and 8 days of starvation for a pair of genotypes. We saw no effect of *hil-1* on reproduction (Fig. S4c). However, for worm length, all three mutants had significantly larger reaction norms to starvation compared to wild type (Figure 3a, b), suggesting HIL-1 supports growth following extended starvation. Because *hil-1(syb9165)* was significantly larger than wild type at day 1 of starvation (actually ∼12 hr), we tested whether this was true in the complete absence of starvation and found no significant effect of *hil-1(syb9165)* on growth (Fig. S4d).

**Figure 3.**
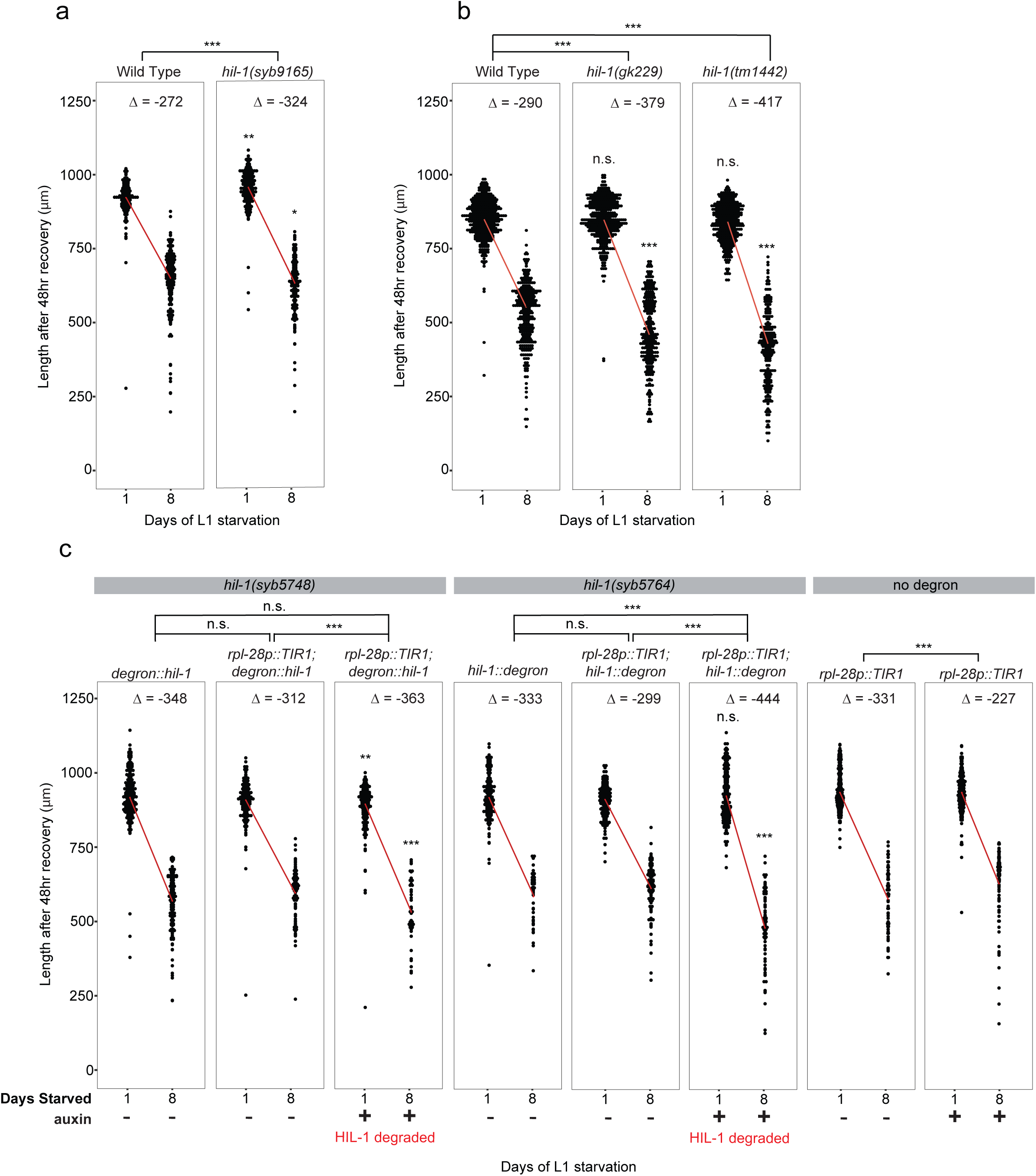
Disruption of *hil-1* exacerbates the effect of extended L1 starvation on larval growth during recovery. Length following 48 hr of recovery from 1 or 8 days L1 starvation is plotted for a) wild type and *hil-1(syb9165),* b) wild type and *hil-1* partial deletion mutants *gk229* and *tm1442*, and c) HIL-1 AID negative control (*degron::hil-1* or *hil-1::degron*) and AID functional depletion strains with and without 100 μM auxin, and a negative control for auxin with *rpl-28p::TIR1*. a-c) Red lines connect the mean values from 1 and 8 days of starvation, plotting the reaction norm for the effect of starvation on size for each genotype. Δ indicates the difference in means between day 1 and day 8 within a genotype (a, b) or genotype + condition (c). Asterisks on the brackets above the panels indicate interaction P-value from a two-factor linear mixed-effect model with genotype and duration of starvation as factors. Asterisks above data points within the panels indicate Tukey-adjusted pairwise comparison within each day of mutant to wild type (a, b) or AID functional depletion strains with auxin compared to no auxin (c). **P < 0.01, ***P < 0.001; 3-4 biological replicates. See also Fig. S4.

Inducible degradation of HIL-1 corroborated the starvation-sensitive phenotype of the *hil-1* deletion mutants. Auxin-inducible degradation (AID) allows for conditional degradation of a protein by tagging it with a degron that is ubiquitinated by transgenic *Arabidopsis* TIR1 when the plant hormone auxin (or an appropriate analog (Martinez et al. 2020)) is added (Zhang et al. 2015). We used *rpl-28p*::*TIR1* to drive strong, ubiquitous expression of TIR1, and we confirmed degradation by depletion of GFP signal (Fig. S4e, f). Degradation was incomplete, suggesting gene function is not null. Nonetheless, degradation of HIL-1 caused a larger effect of extended L1 starvation on length following 48 hr recovery than in controls without degradation for both alleles (Fig. 3c). Adding auxin to the control strain *rpl-28p*::*TIR1* caused a significant decrease in the effect size of extended starvation, suggesting potential off-target effects of TIR1 on starvation resistance. However, the direction of this effect is the opposite of what we see by adding auxin to degrade HIL-1, further supporting the conclusion that HIL-1 supports L1 starvation resistance.

### *hil-1* affects expression of innate immunity genes during starvation

We used bulk RNA-seq to determine how disruption of *hil-1* affects gene expression during L1 starvation. We analyzed wild-type, *hil-1(gk229)*, and *hil-1(tm1442)* worms at ∼12 hr of L1 starvation. The two *hil-1* alleles showed similar gene expression profiles, so we pooled samples from the two alleles as replicates of a single perturbation for further analysis. 463 genes were differentially expressed between *hil-1* and wild type (FDR < 0.1, Fisher’s exact test), with 221 genes upregulated and 242 genes downregulated in the *hil-1* mutants (“repressed by *hil-1*” and “activated by “*hil-1*”, respectively; Fig. 4a, Sup. File 1). Despite *hil-1* presumably functioning as an H1 histone to reduce chromatin accessibility around bound regions, *hil-1* targets displayed no significant chromosomal clustering (Figure S5a, b). We used the Gene Ontology (GO) term enrichment tool GOrilla to investigate the biological function of the genes affected by *hil-1* (Eden et al. 2009). There were no enriched GO-terms for the 221 repressed genes, but the 242 activated genes had 14 enriched GO-terms (FDR < 0.1), most of which are involved in the innate immune and defense response (Fig. 4b, Sup. File 1). We also used CelEsT (Perez 2024) to estimate *hil-1*-dependent changes in transcription factor activity in our RNA-seq experiment (Fig. 4c, Table 2, Sup. File 1). Five transcription factors were estimated to have lower activity in *hil-1* compared to wild-type, and four were estimated to have higher activity. The top three most significant estimations were for decreased activity of bZIP transcription factors CEBP-1, ZIP-4, and ZIP-2, which mediate upregulation of the conserved “Ethanol and Stress Response Element” (ESRE) network in response to exposure to the bacterial pathogen *Pseudomonas aeruginosa* (Estes et al. 2010; McEwan et al. 2016; Tjahjono and Kirienko 2017). These results are consistent with *hil-1* activating immunity genes, as suggested by GO term enrichments (Fig. 4b). We followed up on these observations by comparing the effects of disruption of *hil-1* to exposure to *P. aeruginosa* (Fletcher et al. 2019), and we found a significant overlap between the genes activated by *hil-1* and *P. aeruginosa* PA14 (Fig. 4d). These results suggest that HIL-1 affects transcription of genes that are induced by pathogen exposure and contribute to immunity.

**Figure 4.**
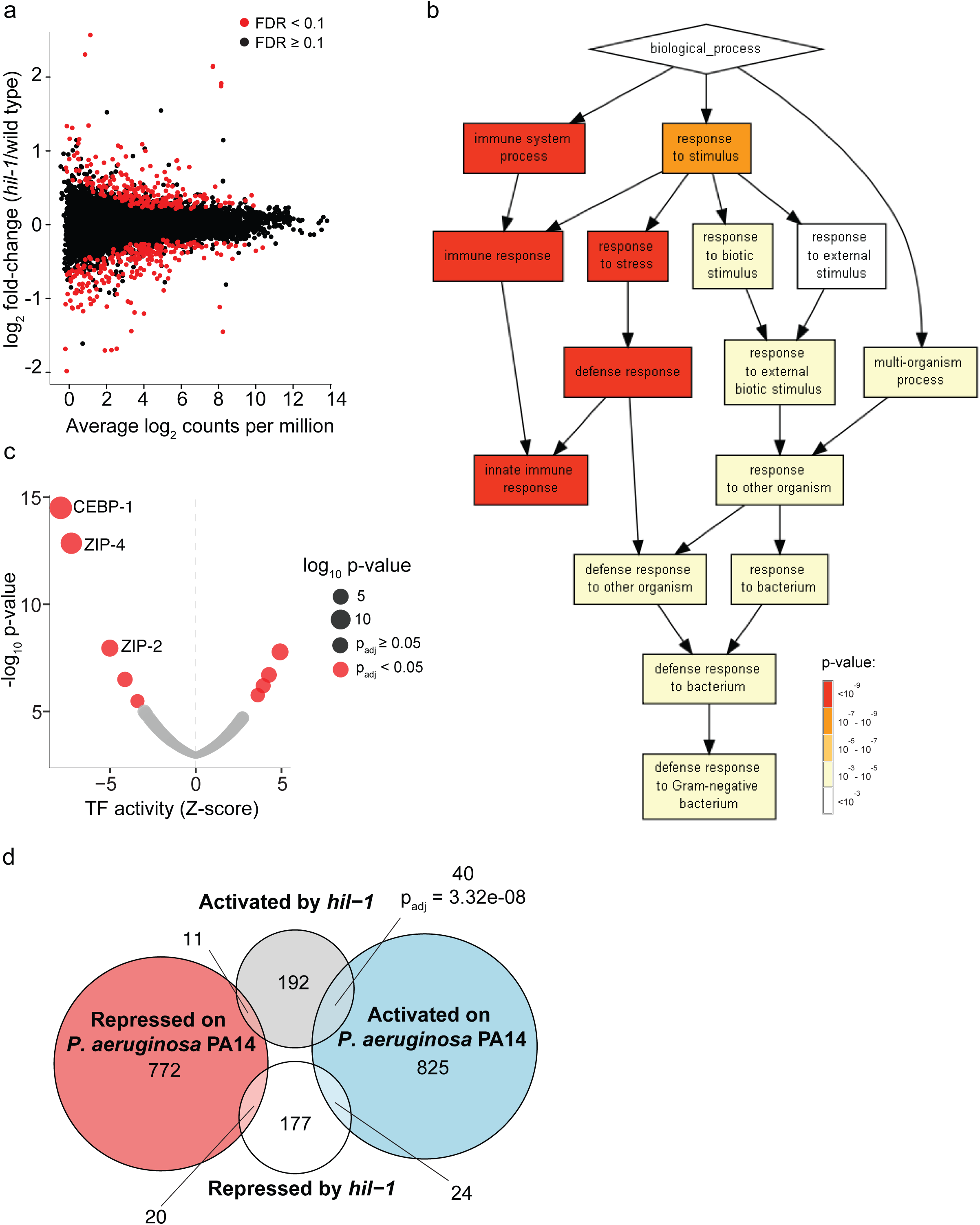
*hil-1* activates genes related to immunity during L1 arrest. a) RNA-seq results of wild type and *hil-1* partial deletion mutants (*gk229* and *tm1442* pooled as replicates) ∼12 hr after hatching without food (starved) with average transcript abundance (counts per million) plotted against log_2_ fold changes between mutant and wild type. Red dots indicate FDR < 0.1 and black dots are insignificant; four biological replicates for wild type and eight for *hil-1*, four per allele pooled. b) GOrilla plot (Eden et al. 2009) for “biological process” GO terms significantly enriched among 242 genes activated by *hil-1* (downregulated in *hil-1* mutant). P-values for enrichment are indicated by color (see legend). c) CelEsT output plotting estimated differences in transcription factor activity given RNA-seq results (*hil-1 /* wild type). Negative values indicate less activity in *hil-1* mutants, and positive values indicate increased activity in *hil-1* mutants. P-values were adjusted for multiple testing (Benjamini-Hochberg). Red dots indicate P < 0.05 and black dots are insignificant. Dot size reflects the absolute value of differences in transcription factor activity (Z-scores). The three most significant transcription factors are labelled. See Table 2 for all significant transcription factors. d) Overlap of genes activated or repressed by *hil-1* and genes differentially expressed on *Pseudomonas aeruginosa* PA14 (Fletcher et al. 2019). Significant adjusted P-values from hypergeometric tests are indicated in the plot; all other P-values were insignificant and not included in the plot. See also Fig. S5.

### *hil-1* promotes survival during exposure to *Pseudomonas aeruginosa*

We exposed wild-type worms and *hil-1* mutants to *P. aeruginosa* PA14 using the “fast-killing” assay (Tan et al. 1999) to determine if *hil-1* promotes pathogen resistance. There was no significant effect on survival for any of the three *hil-1* deletion mutants compared to wild type (Fig. S6). However, we did not see induction of HIL-1 expression on PA14 with our GFP reporters, and it is hardly expressed in larvae fed *E. coli* (Fig. 1a, c; S1; 2c, f; S2), so it is not surprising that disruption of *hil-1* would have no effect under these conditions. Decreasing IIS with a *daf-2* mutation increases resistance to pathogens, including PA14 (Evans et al. 2008; Garsin et al. 2003). We predicted there may be an effect of disruption of *hil-1* in a *daf-2(e1370)* background since *hil-1* expression would be activated in the control (Fig. 2c). *daf-2(e1370)* was very long-lived compared to wild type, as expected, with some individuals surviving over 20 days on “fast-killing” plates (Fig. 5a). All three *hil-1* deletion mutants significantly reduced survival in the *daf-2(e1370)* background, indicating that *hil-1* promotes survival of *P. aeruginosa* exposure in reduced IIS conditions. As a control for survival on *P. aeruginosa*, we tested lifespan on *E. coli* OP50 for the three *hil-1* mutants in the *daf-2(e1370)* background (Fig. 5b, c). Contrary to survival on *P. aeruginosa*, *hil-1(syb9165)* showed a modest but significant increase in survival compared to the control (Fig. 5b), while the partial deletion alleles did not significantly affect lifespan (Fig. 5c). These results support the conclusion that *hil-1* specifically supports survival during exposure to *P. aeruginosa*, though the effect on OP50 lifespan was unexpected.

**Figure 5.**
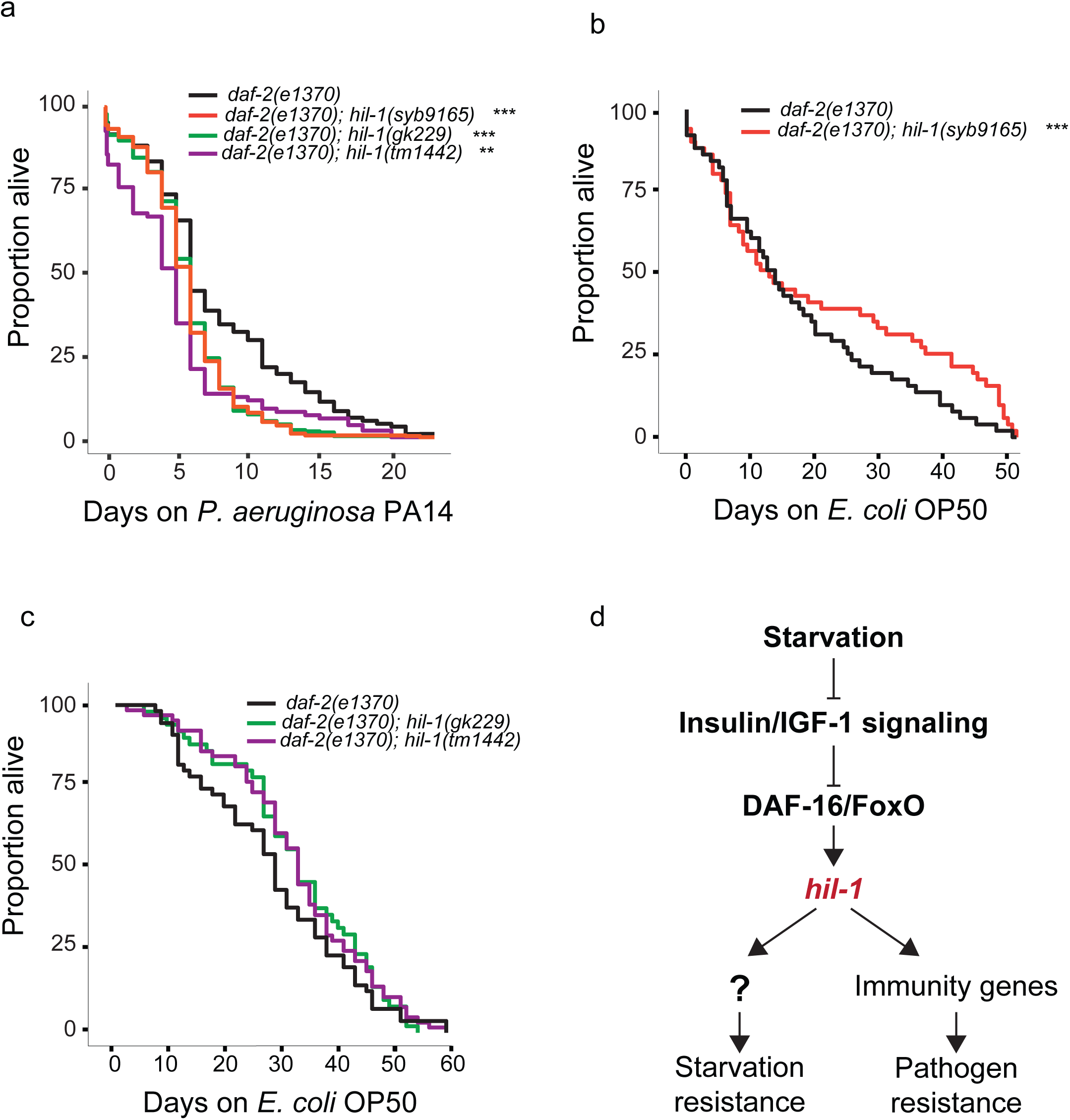
*hil-1* mutants are sensitive to *Pseudomonas aeruginosa* under reduced IIS. a) Survival of *daf-2(e1370)* and *hil-1* mutants in *daf-2(e1370)* background in PA14 fast-killing assay with L4 larvae. b, c) *E. coli* OP50 control for lifespan of *daf-2(e1370)* and *daf-2(e1370); hil-1(syb9165)* (b) or *daf-2(e1370); hil-1(gk229)* and *daf-2(e1370); hil-1(tm1442)* (c). **P < 0.01, ***P < 0.001 (log-rank, Bonferroni-corrected P-value against wild type); 3-4 biological replicates. d) A model for the regulation and function of *hil-1*. DAF-16/FoxO activates *hil-1* transcription when IIS is low, and HIL-1 promotes resistance to starvation and bacterial pathogens (See Discussion). Refer to Fig. S6.

Because *hil-1* affects expression of immunity genes and promotes survival of worms exposed to *P. aeruginosa*, we sought to identify genes that may mediate the effect of *hil-1* on pathogen survival. We identified candidate genes as being 1) activated by *hil-1,* 2) annotated for the GO-term “defense response to other organism”, 3) and upregulated in worms exposed to *P. aeruginosa* PA14 (Fletcher et al. 2019). In addition to the resulting 11 genes, we included *nipi-3/TRIB1*, a Tribbles pseudokinase which regulates CEBP-1 activity to promote resistance to PA14 (McEwan et al. 2016), is activated by *hil-1*, and has overlapping targets with *hil-1* (Table 3), though it was not significantly upregulated on PA14. We cultured *daf-2(e1370)* mutants on RNAi bacteria for each of our 12 candidate genes, with *hil-1* RNAi and empty vector as controls, until they reached the L4 stage, then transferred them to PA14 fast-killing plates to score survival (Fig. S6b, c). Genes were split into two sets, with controls (empty vector and *hil-1*) repeated in multiple replicates for each set. *hil-1* RNAi reduced survival in both sets of assays, but it was only significant in set 2. Lack of a consistent, robust effect of *hil-1* RNAi suggests it will be difficult to detect effects of genes activated by HIL-1. Of the genes tested, *nipi-3, C08E8.4,* and *spp-18* significantly reduced survival, whereas *F08G2.5* unexpectedly increased survival. In conclusion, genes activated by *hil-1* during starvation affect survival of worms exposed to *P. aeruginosa*, consistent with these genes mediating the effect of *hil-1* on pathogen resistance, though we have not demonstrated that their function depends on *hil-1*.

## DISCUSSION

Eukaryotic genomes frequently encode multiple H1 histones, but the extent to which they have distinct vs. overlapping regulation and function is unclear. *hil-1* is unique among the nine annotated H1 variants in *C. elegans* based on its sequence and expression. We characterized the expression of HIL-1 with endogenous GFP reporters, confirming that it is upregulated in starved L1 and dauer larvae and downregulated in response to feeding, and revealing that it is expressed broadly and localized to nuclei. We also show that upregulation of HIL-1 requires *daf-16/FoxO* under the control of IIS. HIL-1 does not have pervasive effects on gene expression, but it activates immunity-related genes, potentially via bZIP transcription factors. Phenotypic analysis reveals that *hil-1* has a modest effect on L1 starvation resistance and that it promotes survival of *daf-2/InsR* mutants exposed to *P. aeruginosa*. This work shows that *hil-1* is unique among *C. elegans* H1 histone genes in its regulation and function, connecting nutrient availability to immunity (Fig. 5d).

### HIL-1 is abundantly expressed in nuclei throughout the animal during starvation

HIL-1 expression has been analyzed in fed worms with antibodies and multicopy reporter genes (Jedrusik et al. 2002), but its expression and function during starvation have not been characterized. HIL-1 expression is most prominent in the pharynx with expression in a variety of additional cell types. Remarkably, Jedrusik *et al*. reported that HIL-1 is localized to the cytoplasm in addition to nuclei, and that it is not localized to chromosomes, where H1 histones are typically observed. Using a pair of endogenous reporter genes (with GFP inserted at the C- and N-termini), we found that HIL-1 expression is hardly detectable in fed L1 larvae, though the pharynx is most prominent. In contrast, HIL-1 expression is dramatically upregulated in starved L1 and dauer larvae, with expression throughout the animal and predominantly nuclear. *hil-1* mRNA and protein levels also decline rapidly in response to feeding starved larvae. We believe our results differ from Jedrusik *et al*. primarily because they did not examine starved larvae, though it is possible that artefacts of antibody staining or multicopy reporters could also be an issue. In addition, we show that HIL-1 expression is sensitive to nutrient availability, with different levels of expression in response to varying nutrient availability, as if HIL-1 functions as a graded nutrient sensor governed by IIS. The HIL-1 reporters generated in this study could be valuable as indicators of nutrient status/IIS activity in *C. elegans*.

### *hil-1* promotes starvation resistance

Dramatic upregulation of HIL-1 during starvation suggests a functional role in acclimation to starvation. However, loss of *hil-1* did not significantly affect L1 starvation survival. Starvation survival is not a particularly sensitive assay, and it could be that we did not have adequate power to observe a significant effect. Extended starvation compromises growth and reproduction upon recovery, and some genotypes and starvation conditions affect recovery without affecting survival (Baugh and Hu 2020).

We also did not see a starvation-dependent effect of *hil-1* on reproduction. However, for growth during recovery from extended starvation, all three deletion alleles of *hil-1* plus both AID alleles revealed an interaction (non-additivity) between *hil-1* and starvation, with larvae growing disproportionately less after extended starvation with disruption of *hil-1*, suggesting HIL-1 promotes starvation resistance. Alternatively, HIL-1 could prime worms during starvation to respond to food, or it could engage additional effectors upon recovery from starvation, resulting in a recovery-specific phenotype. Incomplete degradation of HIL-1 reporter proteins during recovery from starvation could be seen to support the possibility of HIL-1 function during recovery, but decay dynamics differed for the two alleles, and residual GFP signal in larvae recovering from L1 starvation was very low compared to starvation. Furthermore, residual GFP during recovery could result from incomplete degradation and persistence of a fluorescent, non-functional protein fragment. We believe the simplest interpretation of our results is that HIL-1 functions in starved larvae to support growth upon feeding. L1 starvation compromises mitochondrial function (Hibshman et al. 2018), and HIL-1 could support starvation resistance by activating the ESRE network via CEBP-1, ZIP-4, and ZIP-2 (see below) (Tjahjono and Kirienko 2017). In support of this hypothesis, *zip-2* and *cebp-2* mutants are starvation sensitive (Yan et al. 2025).

### *hil-1* promotes pathogen resistance

We report that *hil-1* affects expression of a few hundred genes during L1 starvation, and many of the genes activated by *hil-1* are associated with the innate immune response and bacterial pathogen exposure. We also show that *hil-1* and some of the genes it activates (three out of 12 tested), appear to promote survival of worms exposed to *P. aeruginosa* in a *daf-2/InsR* mutant background (see below). *nipi-3* is one of these three genes, and this Tribbles pseudokinase ortholog is known to be required for activation of intestinal immune genes and survival on *P. aeruginosa* (McEwan et al. 2016), corroborating our results.

Intestinal NIPI-3 negatively regulates the bZIP C/EBP transcription factor CEBP-1 to regulate immunity genes and protect worms against infection (McEwan et al. 2016). In turn, CEBP-1 positively regulates its own expression and *nipi-3* expression (Wu et al. 2021). *nipi-3* mutants display abnormal induction of immunity genes, but pathogen susceptibility of *nipi-3* mutants is rescued when *cebp-1* is mutated (McEwan et al. 2016). In addition to being activated by *nipi-3*, we show that *cebp-1* is activated by *hil-1*, and CEBP-1 had the most significant reduction in estimated transcription factor activity in *hil-1* mutants, followed by two other bZip transcription factors, ZIP-2 and ZIP-4. These three transcription factors upregulate the ESRE defense network in response to mitochondrial stress caused by *P. aeruginosa* exposure (Estes et al. 2010; McEwan et al. 2016; Tjahjono and Kirienko 2017), suggesting that HIL-1 could contribute to mitochondrial surveillance. In further support of CelEst-based transcription factor activity estimates, there is significant overlap between genes activated by *hil-1* and bound by CEBP-1 (Table 3). Curiously, genes repressed by *nipi-3* are enriched among those activated by *hil-1* (Table 3). We suspect this apparent paradox stems from the feedback regulation of NIPI-3 and CEBP-1 (Wu et al. 2021). The PMK-1 p38 MAPK pathway has been reported to function downstream of NIPI-3 and CEBP-1 to affect immunity (Kim et al. 2016; Wu et al. 2021), but it has also been reported to function in parallel (McEwan et al. 2016). There is no significant overlap between genes activated or repressed by *hil-1* and *pmk-1*-dependent genes, suggesting that *hil-1* affects immunity independent of *pmk-1* (Table 3). The activation of innate immunity genes by *hil-1* during starvation, as well as *hil-1* supporting pathogen resistance, suggests that HIL-1 contributes to starvation-dependent transcriptional activation of immunity (Fig. 5d). HIL-1 potentially activates immunity though the NIPI-3/CEBP-1 pathway and the ESRE network, but additional experiments are required to test this hypothesis.

**Table 3.**
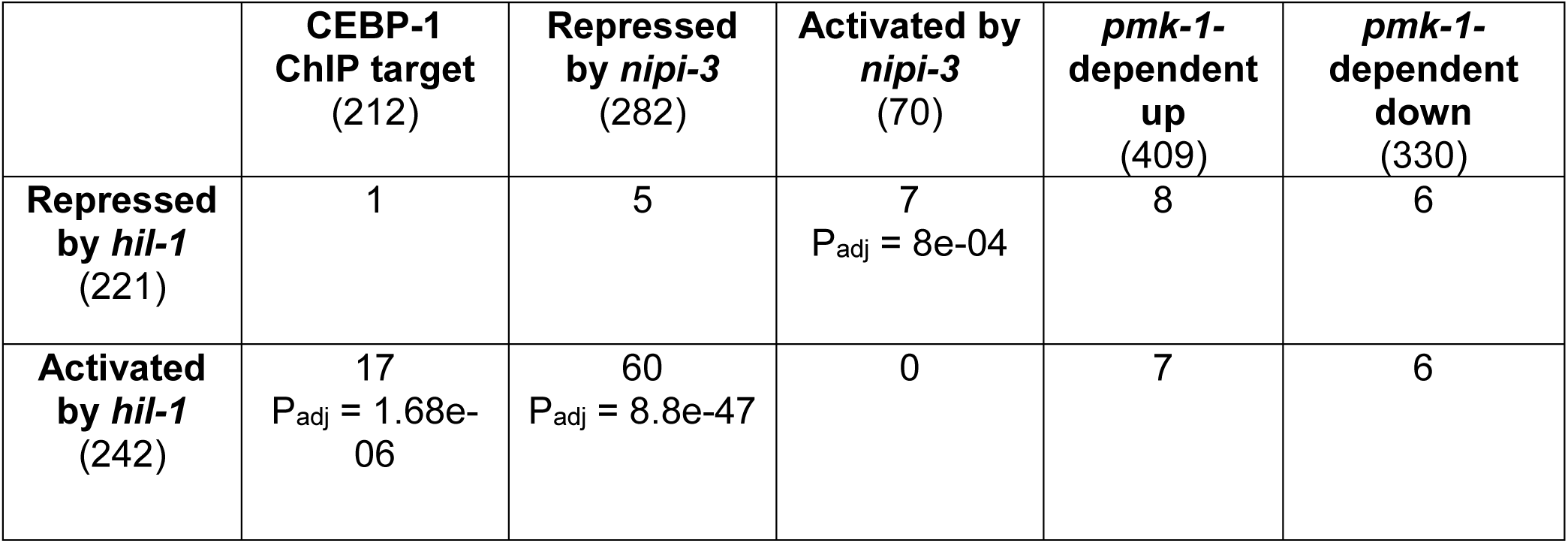
Comparison of *hil-1* targets and components of *nipi-3/TRIB1* immunity pathway. Differentially expressed genes in *hil-1* mutants compared to ChIP targets of CEBP-1 (Kim et al. 2016), differentially expressed genes on PA14 in a *nipi-3(fr4)* mutant (McEwan et al. 2016), and genes that are differentially expressed on PA14 in a *pmk-1*-dependent fashion (Fletcher et al. 2019). The total number of genes in each group is listed in parentheses under group name. The number of genes two groups share are indicated in the table. Only significant adjusted P-values are listed (hypergeometric test on overlap).

Emerging evidence suggests conserved function of linker histones in responding to environmental cues to regulate innate immunity (Li et al. 2022). Another *C. elegans* H1 variant, *his-24*, alters innate immunity gene expression in response to infection to promote survival (Studencka et al. 2012). H1 variants in other systems, such as *Arabidopsis,* have also been tied to immune gene expression following infection to alter survival dynamics (Sheikh et al. 2023).

### Insulin/IGF signaling and pathogen resistance

Reduced IIS and early-life starvation promote pathogen resistance in *C. elegans* (Evans et al. 2008; Falsztyn et al. 2025; Garsin et al. 2003), and mammalian studies have revealed a physiologically significant link between IGF-1 signaling and immune function (Bæk et al. 2024; Smith 2010). We show that IIS regulates expression of HIL-1, and that *hil-1* affects expression of immunity genes and resistance to a bacterial pathogen. However, *hil-1* contributes to pathogen resistance only under reduced IIS. We believe this is due to its negligible expression in fed wild-type worms and induction in *daf-2* mutants. We speculate that this link between nutrient signaling and immune function may reflect evolution with limited benign food sources available to starving worms, making them less selective when emerging from starvation, such that pre-activation of the immune system is adaptive. Alternatively, wild worms may consume relatively small amounts while avoiding bacterial pathogens, and induction of *hil-1* could support fitness in such a scenario. Speculation aside, our results show that *hil-1* contributes to nutrient and IIS-dependent effects on pathogen resistance.

### Conditional regulation of HIL-1 and other histones

All eight detected H1 genes displayed nutrient-dependent regulation in fed and starved L1 larvae. Starvation can change the phosphorylation state of H1 histones in ciliates (Dou et al. 2005), and water deficit can impact expression of specific H1 variants in tomatoes (Scippa et al. 2000). These observations suggest that *C. elegans* H1 histones are not alone in being responsive to environmental conditions. *hil-1* is unique among the *C. elegans* H1 histone genes for being so dramatically upregulated during starvation. *hil-1* is also unique for being directly activated by DAF-16/FoxO. However, *hil-1* expression oscillates with the molting cycle in developing larvae, with upregulation coinciding with lethargus when larvae transiently cease eating and undergo ecdysis {Meeuse, 2020 #170}. Thus, *hil-1* expression is not restricted to outright starvation.

*hil-1* could function redundantly with other H1 variants, particularly *hil-7*, which is also upregulated in starvation. However, HIL-7 lacks a C-terminal domain and therefore is unlikely to function as a canonical H1 histone. To the extent that these paralogs have specific function, it is unclear whether such specificity is due to protein structure or differential expression. Coding sequences could be swapped between H1 genes to address this question. Notably, *hil-2* and *his-24* have the opposite expression pattern of *hil-1*, with *daf-16*-dependent downregulation during starvation, suggesting they could complement *hil-1* function in fed larvae. As precedent, there are 17 members of the H2B family in *C. elegans*, all but one of them (HIS-41) is degraded during L1 starvation, and histone turnover is critical to an appropriate starvation response (Zhu et al. 2023). In addition, chromatin compaction is grossly affected in multiple tissues during L1 starvation, also affecting gene expression (Al-Refaie et al. 2024; Belew et al. 2021). Together these observations suggest that the profound effects of nutrient availability on gene regulation involve extensive changes to chromatin structure, potentially mediated by conditional modulation of histone variant expression.

## DATA AVAILABILITY STATEMENT

All raw data, analysis, and supplementary files can be found at https://github.com/kinseyfish/hil-1.git. The data associated with this manuscript have been deposited in Figshare and will be made publicly available upon publication. Raw and processed data files from RNA-seq can be accessed through the National Center for Biotechnology Gene Expression Omnibus (GEO; http://www.ncbi.nlm.nih.gov/geo) with the accession number <u>GSE318171</u>.

## ACKNOWLEDGEMENTS

This work was funded by the National Institutes of Health (NIGMS awards R01GM117408, R01GM143159, and R35GM156356 to L.R.B.). Some strains were provided by the CGC, which is funded by NIH Office of Research Infrastructure Programs (P40 OD010440). We would like to thank WormBase and the Alliance of Genomes. We would also like to thank Sevinc Ercan for providing the *hil-1(syb9165)* strain, which was created by Suny Biotech and funded by the NIGMS award R35GM130311.

---

**Figure S1. *hil-1* mRNA abundance decreases rapidly upon feeding starved L1 larvae.** Related to Fig. 1. mRNA-seq results (average transcript abundance with 95% confidence intervals) is plotted over time (0 – 6 hr) after feeding L1 larvae that had been starved for ∼12 hr (Maxwell et al. 2012).

**Figure S2. Widespread nuclear expression of HIL-1 with multiple reporters in starved L1 and dauer larvae.** Related to Fig. 2b. 1000x nomarski and GFP images of *hil-1(syb5764)* (a, c) or *hil-1(syb5748*) (b) ∼12 hr after hatching with or without food (fed and starved, respectively) (a) or in fed L3 or starved dauer larvae (b, c). Arrows point to location of GFP expression above background autofluorescence in fed larvae.

**Figure S3. HIL-1 protein expression remains elevated after feeding starved larvae.** Related to Fig. 2e. 1000x Nomarski and GFP images of worms that had residual expression in *hil-1(syb5748)* (a, c) and *hil-1(syb5764*) (b, d) at 12 hr (a, b) and 48 hr (c, d) after hatching with food (continuously fed) or recovery from ∼12 hr L1 starvation (starved then fed). Longer exposure times were used to capture residual expression. Arrows represent locations where GFP is seen above background autofluorescence in continuously fed larvae.

**Figure S4. *hil-1* does not significantly affect L1 starvation survival or reproduction following extended starvation.** Related to Fig. 3. a, b) Survival is plotted over time during L1 arrest for wild type and *hil-1* full deletion mutant (a) or wild type and *hil-1* partial deletion mutants (b). Survival was scored each day (50-150 worms). A curve was fit by logistic regression for plotting averaged data, with individual measurements indicated as points. No significant differences from wild type were detected (pairwise t-test on median survivals from replicates); three (b) or four (a) biological replicates. c) Brood size following 48 hr of recovery from 1 or 8 days L1 starvation for wild type and *hil-1(syb9165).* Red lines connect the mean values from 1 and 8 days of starvation, plotting the reaction norm for the effect of starvation on brood size for each genotype. Δ indicates the difference in means between day 1 and day 8 within a genotype. “n.s.” stands for “not significant; above the panels “n.s.” indicates interaction P-value from a two-factor linear mixed-effect model; “n.s.” above data points within the panels indicates Tukey-adjusted pairwise comparison within each day for mutant to wild type. d) Length of wild type and *hil-1(syb9165)* at 0 d of starvation. P-value from linear mixed-effect model not significant. e) 1000x Nomarski and GFP images with and without addition of auxin analog K-NAA for both *hil-1* AID alleles and a wild-type control (autofluorescence). f) Average background-corrected GFP intensity ∼12 hr after hatching without food (starvation) for both *hil-1* reporters (which also include an AID degron) crossed to *rpl-28p::TIR-1* with H_2_O or 100 µM water-soluble auxin analog K-NAA. ***P < 0.001 (one-way ANOVA). AID causes substantial but incomplete protein degradation.

**Figure S5. *hil-1* targets are not clustered along chromosomes.** Related to Fig. 4. Start position of genes that were significantly down- (a) or upregulated (b) in the *hil-1* mutants in RNA-seq are plotted along chromosomes. Gray bars represent chromosome length. P = n.s. for both sets of genes (permutation test on spatial clustering of target genes using regioneR).

**Figure S6. *hil-1* does not affect pathogen resistance in wild-type background, but its targets contribute to pathogen resistance under reduced IIS. Related to Fig. 5**. a) Survival of wild type and *hil-1* mutants on PA14 fast-killing assay with L4 larvae. None were significantly different from wild type based on the log-rank, Bonferroni-corrected P-value. b, c) PA14 fast-killing assay in *daf-2(e1370)* L4 larvae after feeding RNAi. Empty vector is a negative control. The screen was done in two separate batches (set 1 (b) and set 2 (c)). *P < 0.05, **P < 0.01, ***P < 0.001 (log-rank, Bonferroni-corrected P-value against empty vector control); 4 biological replicates per batch.

## REFERENCES

Al-Refaie N, Padovani F, Hornung J, Pudelko L, Binando F, Del Carmen Fabregat A, Zhao Q, Towbin BD, Cenik ES, Stroustrup N et al. 2024. Fasting shapes chromatin architecture through an mtor/rna pol i axis. Nat Cell Biol. 26(11):1903–1917.

Ashburner M, Ball CA, Blake JA, Botstein D, Butler H, Cherry JM, Davis AP, Dolinski K, Dwight SS, Eppig JT et al. 2000. Gene ontology: Tool for the unification of biology. The gene ontology consortium. Nat Genet. 25(1):25–29.

Bæk O, Rasmussen MB, Gerts T, Aunsholt L, Zachariassen G, Sangild P, Nguyen DN. 2024. Insulin-like growth factor 1 associated with altered immune responses in preterm infants and pigs. Pediatr Res. 95(1):120–128.

Baugh LR. 2013. To grow or not to grow: Nutritional control of development during caenorhabditis elegans l1 arrest. Genetics. 194(3):539–555.

Baugh LR, Demodena J, Sternberg PW. 2009. Rna pol ii accumulates at promoters of growth genes during developmental arrest. Science. 324(5923):92–94.

Baugh LR, Hu PJ. 2020. Starvation responses throughout the caenorhabditiselegans life cycle. Genetics. 216(4):837–878.

Baugh LR, Sternberg PW. 2006. Daf-16/foxo regulates transcription of cki-1/cip/kip and repression of lin-4 during c. Elegans l1 arrest. Curr Biol. 16(8):780–785.

Belew MD, Chien E, Wong M, Michael WM. 2021. A global chromatin compaction pathway that represses germline gene expression during starvation. J Cell Biol. 220(9).

Belfiore A. 2007. The role of insulin receptor isoforms and hybrid insulin/igf-i receptors in human cancer. Curr Pharm Des. 13(7):671–686.

Boucher J, Kleinridders A, Kahn CR. 2014. Insulin receptor signaling in normal and insulin-resistant states. Cold Spring Harb Perspect Biol. 6(1).

Carter ME, Brunet A. 2007. Foxo transcription factors. Current Biology. 17(4):R113–R114.

Castro PV, Khare S, Young BD, Clarke SG. 2012. Caenorhabditis elegans battling starvation stress: Low levels of ethanol prolong lifespan in l1 larvae. PLoS One. 7(1):e29984.

Catez F, Ueda T, Bustin M. 2006. Determinants of histone h1 mobility and chromatin binding in living cells. Nat Struct Mol Biol. 13(4):305–310.

Chen Y, Chen L, Lun Aaron TL, Baldoni Pedro L, Smyth Gordon K. 2025. Edger v4: Powerful differential analysis of sequencing data with expanded functionality and improved support for small counts and larger datasets. Nucleic Acids Research. 53(2).

Consortium TAoGR. 2024a. Updates to the alliance of genome resources central infrastructure. Genetics. 227(1).

Consortium TGO, Aleksander SA, Balhoff J, Carbon S, Cherry JM, Drabkin HJ, Ebert D, Feuermann M, Gaudet P, Harris NL et al. 2023. The gene ontology knowledgebase in 2023. Genetics. 224(1).

Consortium TU. 2024b. Uniprot: The universal protein knowledgebase in 2025. Nucleic Acids Research. 53(D1):D609–D617.

Di Liegro CM, Schiera G, Di Liegro I. 2018. H1.0 linker histone as an epigenetic regulator of cell proliferation and differentiation. Genes (Basel). 9(6).

Dou Y, Song X, Liu Y, Gorovsky MA. 2005. The h1 phosphorylation state regulates expression of cdc2 and other genes in response to starvation in tetrahymena thermophila. Mol Cell Biol. 25(10):3914–3922.

Durinck S, Spellman PT, Birney E, Huber W. 2009. Mapping identifiers for the integration of genomic datasets with the r/bioconductor package biomart. Nature Protocols. 4(8):1184–1191.

Eden E, Navon R, Steinfeld I, Lipson D, Yakhini Z. 2009. Gorilla: A tool for discovery and visualization of enriched go terms in ranked gene lists. BMC Bioinformatics. 10(1):48.

Estes KA, Dunbar TL, Powell JR, Ausubel FM, Troemel ER. 2010. Bzip transcription factor zip-2 mediates an early response to pseudomonas aeruginosa infection in caenorhabditis elegans. Proc Natl Acad Sci U S A. 107(5):2153–2158.

Evans EA, Chen WC, Tan MW. 2008. The daf-2 insulin-like signaling pathway independently regulates aging and immunity in c. Elegans. Aging Cell. 7(6):879–893.

Falsztyn IB, Jordan JM, Chen J, Zhao W, Chitrakar R, Reinke AW, Baugh LR. 2025. Early life starvation and hedgehog-related signaling activate innate immunity downstream of daf-18/pten and lin-35/rb causing developmental pathology in adult c. Elegans. PLoS Genet. 21(12):e1011641.

Fisher K, Chitrakar R, Baugh LR. 2026. Transcriptome- and phenotype-based epistasis analysis in caenorhabditis elegans reveals daf-16/foxo-dependent and independent effects of daf-2/insr in l1 starvation and recovery. G3 (Bethesda).

Fletcher M, Tillman EJ, Butty VL, Levine SS, Kim DH. 2019. Global transcriptional regulation of innate immunity by atf-7 in c. Elegans. PLoS Genet. 15(2):e1007830.

Garsin DA, Villanueva JM, Begun J, Kim DH, Sifri CD, Calderwood SB, Ruvkun G, Ausubel FM. 2003. Long-lived c. Elegans daf-2 mutants are resistant to bacterial pathogens. Science. 300(5627):1921.

Gel B, Díez-Villanueva A, Serra E, Buschbeck M, Peinado MA, Malinverni R. 2016. Regioner: An r/bioconductor package for the association analysis of genomic regions based on permutation tests. Bioinformatics. 32(2):289–291.

Gems D, Sutton AJ, Sundermeyer ML, Albert PS, King KV, Edgley ML, Larsen PL, Riddle DL. 1998. Two pleiotropic classes of daf-2 mutation affect larval arrest, adult behavior, reproduction and longevity in caenorhabditis elegans. Genetics. 150(1):129–155.

Han SK, Lee D, Lee H, Kim D, Son HG, Yang JS, Lee SV, Kim S. 2016. Oasis 2: Online application for survival analysis 2 with features for the analysis of maximal lifespan and healthspan in aging research. Oncotarget. 7(35):56147–56152.

Harshman SW, Young NL, Parthun MR, Freitas MA. 2013. H1 histones: Current perspectives and challenges. Nucleic Acids Research. 41(21):9593–9609.

Hartman PG, Chapman GE, Moss T, Bradbury EM. 1977. Studies on the role and mode of operation of the very-lysine-rich histone h1 in eukaryote chromatin. The three structural regions of the histone h1 molecule. Eur J Biochem. 77(1):45–51.

Hendzel MJ, Lever MA, Crawford E, Th’ng JP. 2004. The c-terminal domain is the primary determinant of histone h1 binding to chromatin in vivo. J Biol Chem. 279(19):20028–20034.

Hibshman JD, Doan AE, Moore BT, Kaplan REW, Hung A, Webster AK, Bhatt DP, Chitrakar R, Hirschey MD, Baugh LR. 2017. Daf-16/foxo promotes gluconeogenesis and trehalose synthesis during starvation to support survival. eLife. 6:e30057.

Hibshman JD, Leuthner TC, Shoben C, Mello DF, Sherwood DR, Meyer JN, Baugh LR. 2018. Nonselective autophagy reduces mitochondrial content during starvation in caenorhabditis elegans. Am J Physiol Cell Physiol. 315(6):C781–c792.

Hibshman JD, Webster AK, Baugh LR. 2021. Liquid-culture protocols for synchronous starvation, growth, dauer formation, and dietary restriction of caenorhabditis elegans. STAR Protocols. 2(1):100276.

Jedrusik MA, Schulze E. 2001. A single histone h1 isoform (h1.1) is essential for chromatin silencing and germline development in caenorhabditis elegans. Development. 128(7):1069–1080.

Jedrusik MA, Vogt S, Claus P, Schulze E. 2002. A novel linker histone-like protein is associated with cytoplasmic filaments in caenorhabditis elegans. J Cell Sci. 115(Pt 14):2881–2891.

Kalashnikova AA, Rogge RA, Hansen JC. 2016. Linker histone h1 and protein–protein interactions. Biochimica et Biophysica Acta (BBA) - Gene Regulatory Mechanisms. 1859(3):455–461.

Kalashnikova AA, Winkler DD, McBryant SJ, Henderson RK, Herman JA, DeLuca JG, Luger K, Prenni JE, Hansen JC. 2013. Linker histone h1.0 interacts with an extensive network of proteins found in the nucleolus. Nucleic Acids Res. 41(7):4026–4035.

Kaplan RE, Chen Y, Moore BT, Jordan JM, Maxwell CS, Schindler AJ, Baugh LR. 2015. Dbl-1/tgf-β and daf-12/nhr signaling mediate cell-nonautonomous effects of daf-16/foxo on starvation-induced developmental arrest. PLoS Genet. 11(12):e1005731.

Kenyon C. 2011. The first long-lived mutants: Discovery of the insulin/igf-1 pathway for ageing. Philos Trans R Soc Lond B Biol Sci. 366(1561):9–16.

Kim KW, Thakur N, Piggott CA, Omi S, Polanowska J, Jin Y, Pujol N. 2016. Coordinated inhibition of c/ebp by tribbles in multiple tissues is essential for caenorhabditis elegans development. BMC Biol. 14(1):104.

Kirienko NV, Cezairliyan BO, Ausubel FM, Powell JR. 2014. Pseudomonas aeruginosa pa14 pathogenesis in caenorhabditis elegans. Methods Mol Biol. 1149:653–669.

Kolberg L, Raudvere U, Kuzmin I, Adler P, Vilo J, Peterson H. 2023. G:Profiler—interoperable web service for functional enrichment analysis and gene identifier mapping (2023 update). Nucleic Acids Research. 51(W1):W207–W212.

Langmead B, Trapnell C, Pop M, Salzberg SL. 2009. Ultrafast and memory-efficient alignment of short DNA sequences to the human genome. Genome Biol. 10(3):R25.

Lawrence M, Huber W, Pagès H, Aboyoun P, Carlson M, Gentleman R, Morgan MT, Carey VJ. 2013. Software for computing and annotating genomic ranges. PLoS Comput Biol. 9(8):e1003118.

Lee RY, Hench J, Ruvkun G. 2001. Regulation of c. Elegans daf-16 and its human ortholog fkhrl1 by the daf-2 insulin-like signaling pathway. Curr Biol. 11(24):1950–1957.

Li X, Ye Y, Peng K, Zeng Z, Chen L, Zeng Y. 2022. Histones: The critical players in innate immunity. Frontiers in Immunology. Volume 13 - 2022.

Love MI, Huber W, Anders S. 2014. Moderated estimation of fold change and dispersion for rna-seq data with deseq2. Genome Biol. 15(12):550.

Martinez MAQ, Kinney BA, Medwig-Kinney TN, Ashley G, Ragle JM, Johnson L, Aguilera J, Hammell CM, Ward JD, Matus DQ. 2020. Rapid degradation of caenorhabditis elegans proteins at single-cell resolution with a synthetic auxin. G3 (Bethesda). 10(1):267–280.

Maxwell CS, Antoshechkin I, Kurhanewicz N, Belsky JA, Baugh LR. 2012. Nutritional control of mrna isoform expression during developmental arrest and recovery in c. Elegans. Genome Res. 22(10):1920–1929.

McEwan DL, Feinbaum RL, Stroustrup N, Haas W, Conery AL, Anselmo A, Sadreyev R, Ausubel FM. 2016. Tribbles ortholog nipi-3 and bzip transcription factor cebp-1 regulate a caenorhabditis elegans intestinal immune surveillance pathway. BMC Biol. 14(1):105.

Moore BT, Jordan JM, Baugh LR. 2013. Wormsizer: High-throughput analysis of nematode size and shape. PLoS One. 8(2):e57142.

Muñoz MJ, Riddle DL. 2003. Positive selection of caenorhabditis elegans mutants with increased stress resistance and longevity. Genetics. 163(1):171–180.

Murphy CT, Hu PJ. 2013. Insulin/insulin-like growth factor signaling in c. Elegans. WormBook.1–43.

Orrego M, Ponte I, Roque A, Buschati N, Mora X, Suau P. 2007. Differential affinity of mammalian histone h1 somatic subtypes for DNA and chromatin. BMC Biol. 5:22.

Perez MF. 2024. Celest: A unified gene regulatory network for estimating transcription factor activities in c. Elegans. Genetics. 229(3).

Preibisch S, Saalfeld S, Tomancak P. 2009. Globally optimal stitching of tiled 3d microscopic image acquisitions. Bioinformatics. 25(11):1463–1465.

Putri GH, Anders S, Pyl PT, Pimanda JE, Zanini F. 2022. Analysing high-throughput sequencing data in python with htseq 2.0. Bioinformatics. 38(10):2943–2945.

Sanicola M, Ward S, Childs G, Emmons SW. 1990. Identification of a caenorhabditis elegans histone h1 gene family. Characterization of a family member containing an intron and encoding a poly(a)+ mrna. J Mol Biol. 212(2):259–268.

Schindelin J, Arganda-Carreras I, Frise E, Kaynig V, Longair M, Pietzsch T, Preibisch S, Rueden C, Saalfeld S, Schmid B et al. 2012. Fiji: An open-source platform for biological-image analysis. Nature Methods. 9(7):676–682.

Schulenburg H, Félix MA. 2017. The natural biotic environment of caenorhabditis elegans. Genetics. 206(1):55–86.

Schuster E, McElwee JJ, Tullet JM, Doonan R, Matthijssens F, Reece-Hoyes JS, Hope IA, Vanfleteren JR, Thornton JM, Gems D. 2010. Damid in c. Elegans reveals longevity-associated targets of daf-16/foxo. Mol Syst Biol. 6:399.

Scippa GS, Griffiths A, Chiatante D, Bray EA. 2000. The h1 histone variant of tomato, h1-s, is targeted to the nucleus and accumulates in chromatin in response to water-deficit stress. Planta. 211(2):173–181.

Sheikh AH, Nawaz K, Tabassum N, Almeida-Trapp M, Mariappan KG, Alhoraibi H, Rayapuram N, Aranda M, Groth M, Hirt H. 2023. Linker histone h1 modulates defense priming and immunity in plants. Nucleic Acids Res. 51(9):4252–4265.

Sievers F, Wilm A, Dineen D, Gibson TJ, Karplus K, Li W, Lopez R, McWilliam H, Remmert M, Söding J et al. 2011. Fast, scalable generation of high-quality protein multiple sequence alignments using clustal omega. Mol Syst Biol. 7:539.

Smith TJ. 2010. Insulin-like growth factor-i regulation of immune function: A potential therapeutic target in autoimmune diseases? Pharmacol Rev. 62(2):199–236.

Sridhar A, Orozco M, Collepardo-Guevara R. 2020. Protein disorder-to-order transition enhances the nucleosome-binding affinity of h1. Nucleic Acids Res. 48(10):5318–5331.

Studencka M, Konzer A, Moneron G, Wenzel D, Opitz L, Salinas-Riester G, Bedet C, Krüger M, Hell SW, Wisniewski JR et al. 2012. Novel roles of caenorhabditis elegans heterochromatin protein hp1 and linker histone in the regulation of innate immune gene expression. Mol Cell Biol. 32(2):251–265.

Tan MW, Mahajan-Miklos S, Ausubel FM. 1999. Killing of caenorhabditis elegans by pseudomonas aeruginosa used to model mammalian bacterial pathogenesis. Proc Natl Acad Sci U S A. 96(2):715–720.

Tepper RG, Ashraf J, Kaletsky R, Kleemann G, Murphy CT, Bussemaker HJ. 2013. Pqm-1 complements daf-16 as a key transcriptional regulator of daf-2-mediated development and longevity. Cell. 154(3):676–690.

Thomas PD, Ebert D, Muruganujan A, Mushayahama T, Albou L-P, Mi H. 2022. Panther: Making genome-scale phylogenetics accessible to all. Protein Science. 31(1):8–22.

Tjahjono E, Kirienko NV. 2017. A conserved mitochondrial surveillance pathway is required for defense against pseudomonas aeruginosa. PLoS Genet. 13(6):e1006876.

Wu C, Karakuzu O, Garsin DA. 2021. Tribbles pseudokinase nipi-3 regulates intestinal immunity in caenorhabditis elegans by controlling skn-1/nrf activity. Cell Rep. 36(7):109529.

Yan J, Bhanshali F, Shuzenji C, Mendenhall TT, Taylor SKB, Ermakova G, Cheng X, Bai P, Diwan G, Seraj D et al. 2025. Eukaryotic elongation factor 2 kinase efk-1/eef2k promotes starvation resistance by preventing oxidative damage in c. Elegans. Nature Communications. 16(1):1752.

Zhang L, Ward JD, Cheng Z, Dernburg AF. 2015. The auxin-inducible degradation (aid) system enables versatile conditional protein depletion in c. Elegans. Development. 142(24):4374–4384.

Zhu Z, Li D, Jia Z, Zhang W, Chen Y, Zhao R, Zhang YP, Zhang WH, Deng H, Li Y et al. 2023. Global histone h2b degradation regulates insulin/igf signaling-mediated nutrient stress. The EMBO Journal. 42(19):EMBJ2022113328.

